# Purification and characterization of recombinant Rtt109, a fungus-specific histone acetyltransferase, from *Candida albicans*

**DOI:** 10.64898/2026.07.30.741489

**Authors:** Simran Sharma, Vijayan Ramachandran, Rohini Muthuswami, Samudrala Gourinath, Sneha Sudha Komath

## Abstract

Epigenetic regulation of chromatin dynamics via histone acetylation is one of several mechanisms by which eukaryotes regulate gene expression, DNA replication and repair, and maintain genome stability. This function is performed by histone acetyltransferases (HATs). Rtt109 is one such cytoplasmically localized HAT required for H3K56 acetylation found exclusively in fungi. Using recombinantly expressed *Candida albicans* Rtt109 and its chaperones, Vps75 and Asf1, we show that it can acetylate a 20-residue N-terminal H3 peptide in a coupled HAT assay only in the presence of Vps75, but not in the presence of Asf1 *in vitro*. This appears to be due to the fact that Rtt109-Vps75 is a high affinity stable complex, as estimated by biolayer interferometry (BLI) and gel filtration studies. The HAT activity of the Rtt109-Vps75 complex necessarily requires a flexible 118-160 residue loop of Rtt109 but not the C-terminal domain of Vps75. These results are comparable with what has been observed for the *Saccharomyces cerevisiae* Rtt109 homolog. *In silico* screening of 1,350,000 molecules from Life Chemicals Databases identified some likely inhibitors of *C. albicans* Rtt109 and six of them tested for binding to Rtt109 using BLI. The best ligand, F2368-0266, was used to study its effect on steady state enzyme kinetics, and found to be a competitive inhibitor of the peptide substrate but not of acetyl-CoA. Given the importance of Rtt109 in regulating virulence attributes such as hyphal morphogenesis and GPI biosynthesis in *Candida albicans*, and its effect on fungal pathogenesis, these results have significant clinical implications.

## Introduction

Rtt109 was first discovered in *Saccharomyces cerevisiae* as a protein required for histone H3K56 acetylation [1], a modification that we now know is required not only for transcription, but also for DNA replication, repair, chromatin maturation, genomic stability, cell cycle progression and sensitivity to genotoxic agents [2–6]. Its structural homologs, CREB-binding protein (CBP)/p300, play similar roles in metazoans and are implicated in human cancers [7]. However, at the primary sequence level, Rtt109 shows no significant degree of conservation with respect to the CBP/p300 proteins. Indeed, unlike the latter, Rtt109 does not even appear to possess a canonical acetyltransferase domain. It only weakly acetylates histones on its own but when present in complex with either of the histone chaperones, Vps75 or Asf1, it can acetylate non-nucleosomal H3K56 [4,8,9]. The complexes formed by Rtt109 with Vps75 and Asf1 are distinct. Rtt109-Vps75 has been shown to acetylate H3K9 and H3K56 in vitro, and to acetylate H3K9, H3K23, H3K27 and, to a lesser extent, H3K56 in vivo. On the other hand, Rtt109-Asf1 primarily catalyzes H3K56 acetylation on the H3-H4 dimer both in vitro and in vivo [1,10–12]. Given the importance of H3K56 acetylation for genotoxic stress survival, it follows, therefore, that *S. cerevisiae* strains incapable of *RTT109* (*rtt109*Δ) or *ASF1* (*asf1*Δ) expression are sensitive to genotoxic agents [4,9].

The loss of Rtt109 also makes the opportunistic human pathogenic fungus, *C. albicans*, sensitive to genotoxic and oxidative stress and hinders its efficient white to opaque transition [5,13,14]. The protein plays a major role in the expression of GPI anchored proteins on the fungal cell surface as well as in its filamentation and response to antifungal drugs [15–17]. Not surprisingly, then, its loss also attenuates the pathogenesis of the organism in mouse model studies [14]. Thus, it is considered a potential drug target.

Biochemical characterization of *C. albicans* Rtt109 (CaRtt109), which shares nearly 30% sequence identity with *S. cerevisiae* Rtt109 (ScRtt109), is still lacking and is the main focus of the present study. We show that recombinant CaRtt109 interacts better with CaVps75 than with CaAsf1. We also established a coupled HAT assay using a short N-terminal H3 peptide, and show that CaRtt109-CaVps75 but not CaRtt109-CaAsf1 catalyzes acetylation of the substrate. Using this we obtained steady state kinetic parameters for CaVps75-catalyzed CaRtt109 HAT activity. *In silico* screening of a library of small biologically active molecules helped identify potential inhibitors, six of which were studied for their interaction with Rtt109 using biolayer interferometry (BLI). The molecule with the highest affinity in BLI showed moderate inhibition in the HAT assay. Analysis of the data, suggests that the inhibitor specifically competed with the H3 peptide substrate and not with acetyl-CoA.

## Results

### The primary sequence and fold structure of *C. albicans* Rtt109 is similar to *S. cerevisiae* Rtt109

The protein sequences of Rtt109 from *C. albicans* (CaRtt109), *S. cerevisiae* (ScRtt109), *A. fumigatus* (AfRtt109), and *P. carinii* (PcRtt109) were aligned using Clustal Omega [18] and represented using ESPript 3.0 (https://espript.ibcp.fr/ESPript/ESPript/) (Figure 1). It is seen that CaRtt109 (359 amino acid residues) shares ∼30% sequence identity with ScRtt109 (436 amino acid residues) and about 28% with both AfRtt109 (543 amino acid residues) and PcRtt109 (375 amino acid residues). The flexible 130-179 loop region of *Sc*Rtt109 which is required for interaction with Vps75 is present in CaRtt109 (118-160 amino acids), but is missing in AfRtt109 and PcRtt109. It is possible that this is not a conserved feature of fungal Rtt109. Even between CaRtt109 and ScRtt109, we see only ∼14% conservation within this region. In contrast, the key catalytic residues Asp89, Tyr199 and Trp222 identified in ScRtt109 are conserved among all these species [19–21]. Another residue, Asp288 which is required for efficient activity of ScRtt109 [22] is also conserved among the Rtt109 homologs.

**Figure 1.**
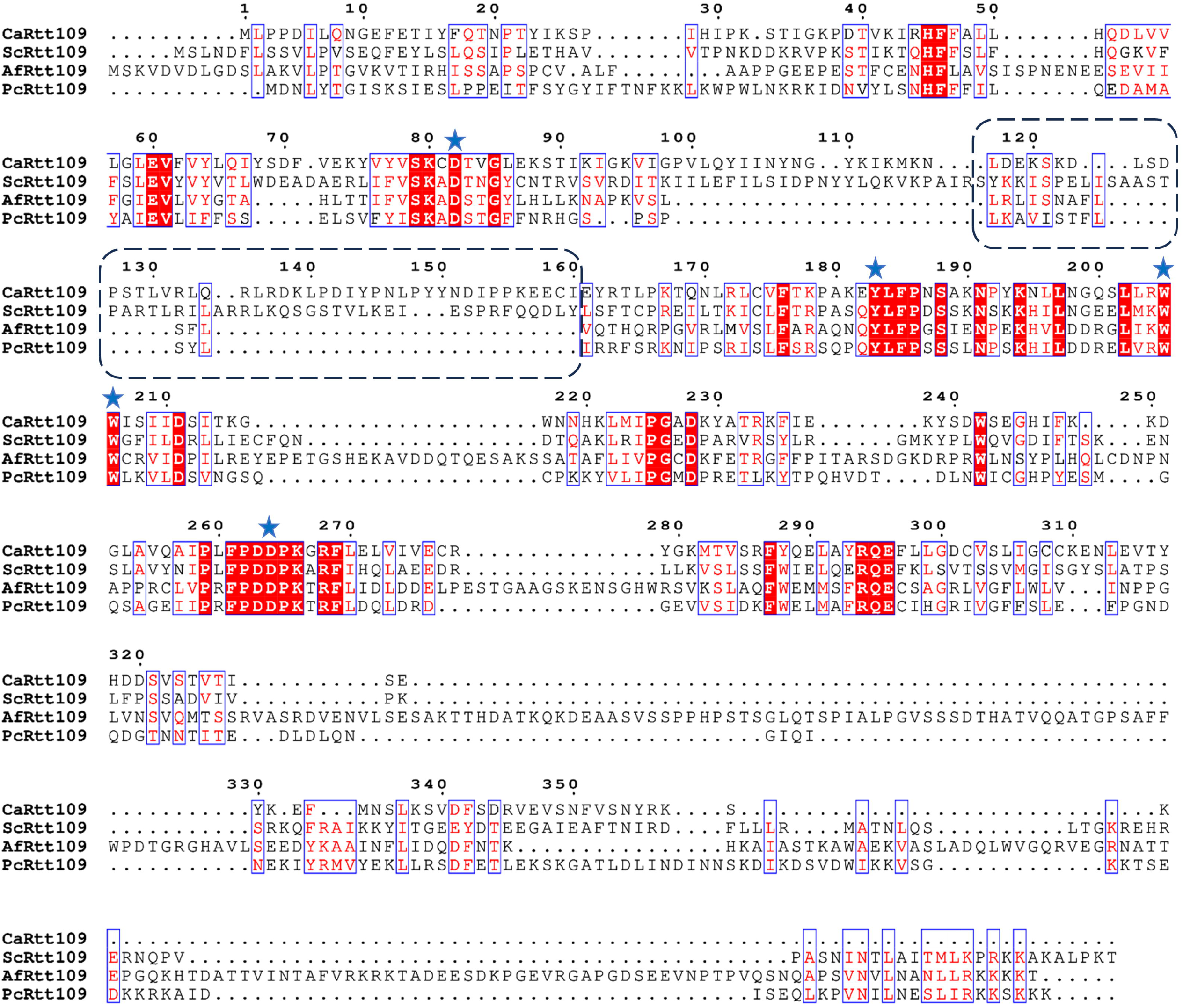
Multiple-sequence alignment of Rtt109 from different species. Multiple-sequence alignment was performed for Rtt109 proteins from *Candida albicans* (CaRtt109; UniProt accession Q5AAJ8), *Saccharomyces cerevisiae* (ScRtt109; UniProt accession Q07794), *Aspergillus fumigatus* (AfRtt109; UniProt accession Q4WUS9), and *Pneumocystis carinii* (PcRtt109; UniProt accession D3G9N3) using Clustal Omega, and the alignment was visualized with ESPript 3.0. Conserved residues are indicated by a red background, whereas similar residues are shown as red color within blue boxes. The region corresponding to the variable loop of Rtt109 (approximately amino acids 118-160) is highlighted by a black dashed box. Residues marked with blue stars denote the catalytic residues identified in ScRtt109 (D89, Y199, W221, W222, and D288), all of which are conserved across the other fungal Rtt109 homologs included in the alignment.

## Cloning, expression and purification of CaRtt109 and its deletion mutant

His_6_-CaRtt109 and His_6_-CaRtt109Δ_Loop_ were cloned in pCold I vector and confirmed by restriction digestion using XhoI and BamHI enzymes (Figure S1 A-B). As explained in the Methods section, His_6_-CaRtt109 and CaHis_6_-Rtt109Δ_Loop_ were individually expressed in Rosetta(DE3) and purified using Ni^+2^-NTA affinity chromatography followed by gel filtration chromatography (GFC) (Figure 2 A-B and Figure 2 C-D, respectively). The GFC HiLoad 16/60 Superdex 200 column used was calibrated using protein standards (Figure S2 A). The apparent molecular weight of both the recombinant proteins corresponded to their expected monomeric size in GFC.

**Figure 2.**
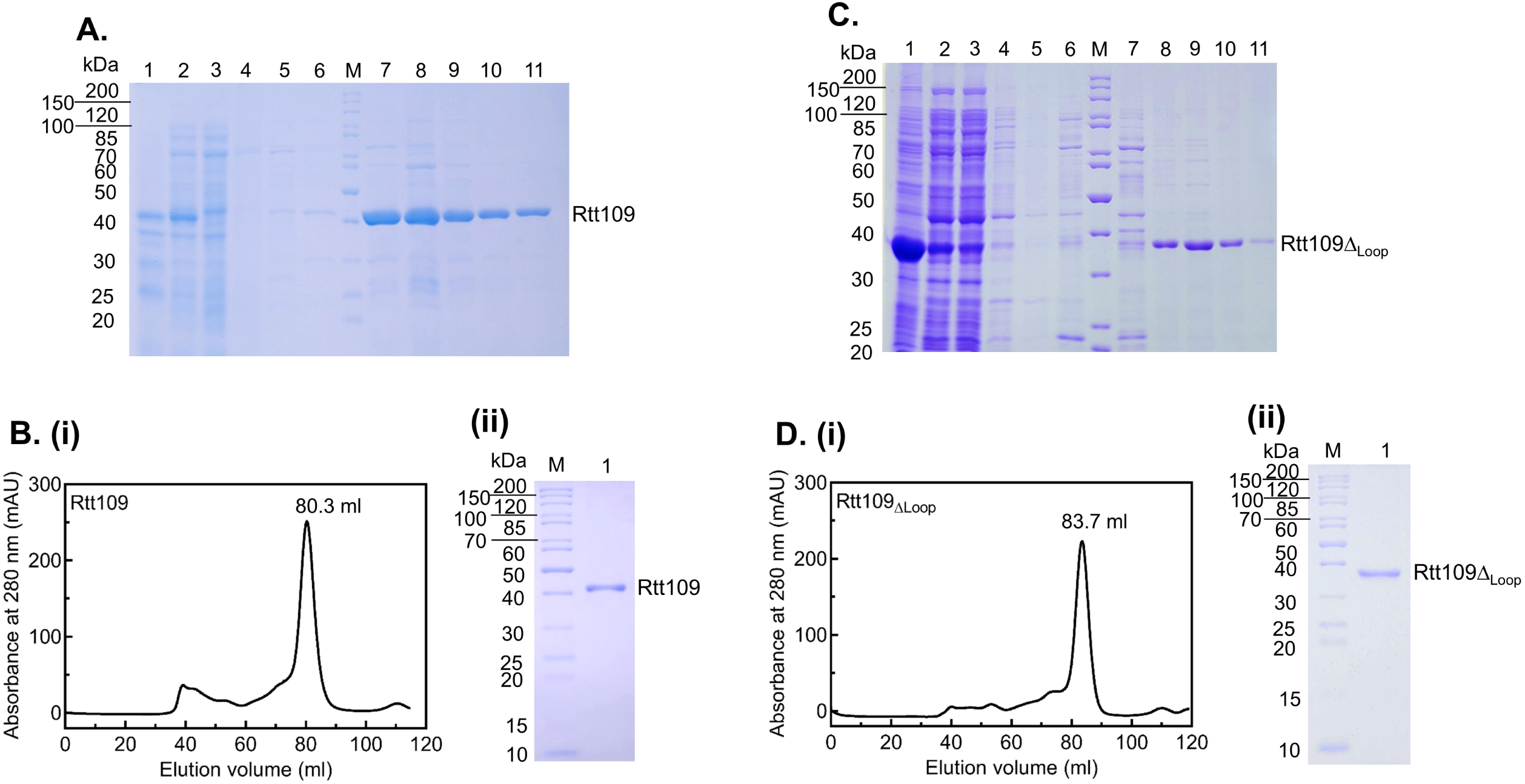
Purification of CaRtt109 and CaRtt109Δ_Loop_. *E. coli* Rosetta(DE3) cells transformed with the respective recombinant plasmids were grown until OD_600nm_ ∼0.6 and protein expression induced using 0.5 mM IPTG at 16 °C for 16 h. The cells were harvested and lysed in 50 mM HEPES-KOH, pH 7.5, containing 300 mM NaCl, 0.5 mM DTT, 5% glycerol, and 1 mM PMSF. **A. SDS-PAGE analysis of the expression and purification of CaRtt109**. Samples from different steps of the Ni^+2^-NTA affinity purification of CaRtt109 were analyzed using 12% SDS-PAGE. Lanes 1 and 2: Pellet and supernatant fractions, respectively, obtained after centrifugation of the cell lysate; Lane 3: Flow-through fraction from the Ni^+2^-NTA column, showing the unbound proteins; Lanes 4-5: wash fractions with buffer containing 300 mM NaCl and 500 mM NaCl, respectively; Lane 6: wash fraction with buffer containing 300 mM NaCl and 10 mM imidazole; Lane M: Molecular weight markers; Lanes 7-11: protein fractions (2 ml each) eluted in the presence of 200 mM imidazole. **B. GFC analysis of purified CaRtt109.** The eluted fractions (7-11) were pooled and concentrated and loaded onto a HiLoad 16/60 Superdex 200 column. **(i)** GFC profile for CaRtt109 shows one peak at 80.3 ml volume, corresponding to the expected M_r_ of a monomer (44 kDa). **(ii)** SDS-PAGE analysis of the peak obtained at 80.3 ml in GFC. Lane M: Molecular weight markers; Lane 1: GFC peak fraction. **C. SDS-PAGE analysis of the expression and purification of CaRtt109Δ_Loop_**. Samples from different steps of the Ni^2+^-NTA affinity purification of CaRtt109Δ_Loop_ were analyzed using 12% SDS-PAGE. Lanes 1 and 2: Pellet and supernatant fractions, respectively, obtained after centrifugation of the cell lysate; Lane 3: Flow-through fraction from the Ni^+2^-NTA column, showing the unbound proteins; Lanes 4-5: wash fractions with buffer containing 300 mM NaCl and 500 mM NaCl, respectively; Lane 6: wash fraction with buffer containing 300 mM NaCl and 10 mM imidazole; Lane M: Molecular weight markers; Lanes 7-11: protein fractions (2 ml each) eluted in the presence of 200 mM imidazole. **D. GFC analysis of purified CaRtt109Δ_Loop_.** The eluted fractions (8-11) were pooled and concentrated and loaded onto a HiLoad 16/60 Superdex 200 column. **(i)** GFC profile for CaRtt109Δ_Loop_ shows one peak at 83.7 ml volume from HiLoad 16/60 Superdex 200 column, which corresponds to the expected M_r_ of a monomer (38 kDa) and **(ii)** SDS-PAGE analysis of the peak obtained at 83.7 ml from GFC. Lane M: Molecular weight markers; Lane 1: GFC peak fraction.

## Cloning, expression and purification of CaVps75 and its deletion mutant

His_6_-CaVps75 and His_6_-CaVps75Δ_C_ (lacking the less-ordered acidic C-terminal tail) were cloned in MCS-I of pRSFDuet1 vector and confirmed by restriction digestion using BamHI and SalI enzymes (Figure S1 C-D). His_6_-CaVps75 and His_6_-CaVps75Δ_C_ were expressed in Rosetta(DE3) and purified using Ni^+2^-NTA affinity chromatography at different NaCl concentrations and then followed by GFC (Figure 3 A-C and Figure 3 D-F, respectively).

**Figure 3.**
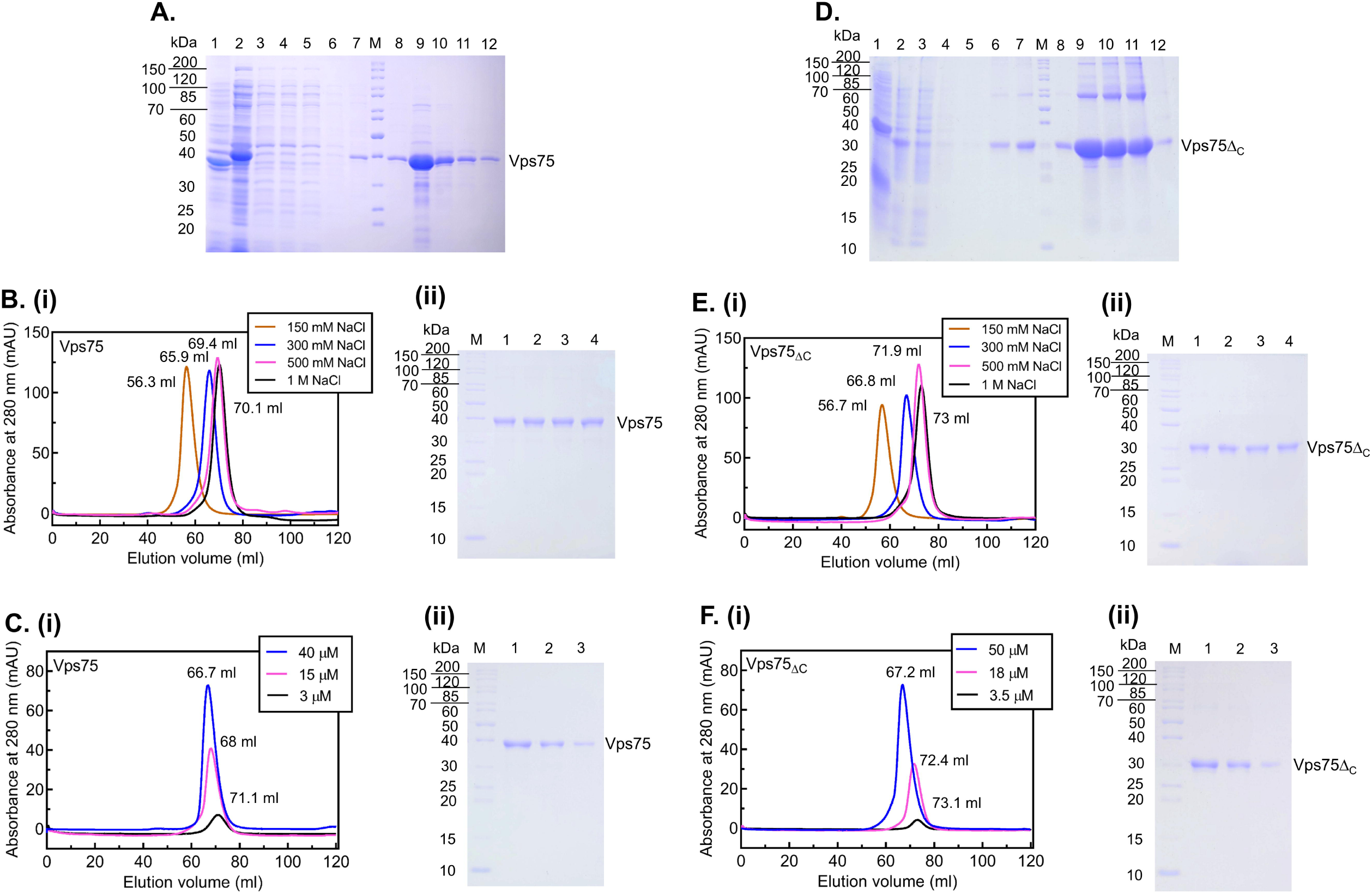
Effect of salt and protein concentration on CaVps75 and CaVps75Δ_C_ oligomerization status. *E. coli* Rosetta(DE3) cells transformed with the respective recombinant plasmids were grown until OD_600nm_ ∼0.6 and protein expression induced using 0.5 mM IPTG at 16 °C for 16 h. The cells were harvested and lysed in 50 mM HEPES-KOH, pH 7.5, containing 300 mM NaCl, 0.5 mM DTT, 5% glycerol, and 1 mM PMSF. **A. SDS-PAGE analysis of the expression and purification of CaVps75**. Lanes 1 and 2: Pellet and supernatant fractions, respectively, obtained after centrifugation of the cell lysate; Lane 3: Flow-through fraction from the Ni^+2^-NTA column, showing the unbound proteins; Lanes 4-5: wash fractions with buffer containing 300 mM NaCl and 500 mM NaCl, respectively; Lanes 6-7: wash fractions with buffer containing 300 mM NaCl with 10 mM and 20 mM imidazole, respectively; Lane M: Molecular weight markers; Lanes 8-12: eluted samples collected in 2 ml fractions at 200 mM imidazole. **B. Ionic strength dependent oligomerization of CaVps75. (i)** Gel filtration chromatography (GFC) profile for CaVps75 eluted from HiLoad 16/60 Superdex 200 column at 56.3 ml, 65.9 ml, 69.4 ml and 70.1 ml, which correspond to M_r_ of 295.1 kDa, 131 kDa, 97.7 kDa, and 91.2 kDa at 150 mM (brown curve), 300 mM (blue curve), 500 mM (pink curve), and 1 M NaCl (black curve), respectively. **(ii)** SDS-PAGE analysis of elute fraction corresponding to each of the GFC peaks. Lane M: Molecular weight markers; Lane 1-4: Elute fractions corresponding to the peaks from GFC at 150 mM, 300 mM, 500 mM, and 1 M NaCl, respectively. **C. Protein concentration dependent oligomerization of CaVps75. (i)** GFC profile for CaVps75 eluted from HiLoad 16/60 Superdex 200 column at 66.7 ml, 68 ml, and 71.1 ml, which correspond to M_r_ of 123 kDa, 109.6 kDa, and 85.1 kDa at 40 μM (blue curve), 15 μM (pink curve), and 3 μM (black curve) protein, respectively. **(ii)** SDS-PAGE analysis of elute fractions corresponding to each of the GFC peaks. Lane M: Molecular weight markers; Lane 1-3: Elute fractions corresponding to the peaks from GFC at 40 μM, 15 μM, and 3 μM protein, respectively. **D. SDS-PAGE analysis of the expression and purification of CaVps75Δ_C_**. Lanes 1 and 2: Pellet and supernatant fractions, respectively, obtained after centrifugation of the cell lysate; Lane 3: Flow-through fraction from the Ni^+2^-NTA column, showing the unbound proteins; Lanes 4-5: wash fractions with buffer containing 300 mM NaCl and 500 mM NaCl, respectively; Lanes 6-7: wash fractions with buffer containing 300 mM NaCl with 10 mM and 20 mM imidazole, respectively; Lane M: Molecular weight markers; Lanes 8-12: eluted samples collected in 2 ml fractions at 200 mM imidazole. **E. Ionic strength dependent oligomerization of CaVps75Δ_C_. (i)** GFC profile for Vps75Δ_C_ eluted from HiLoad 16/60 Superdex 200 column at 56.7 ml, 66.8 ml, 71.9 ml, and 73ml which correspond to M_r_ of 281.8 kDa, 120.2 kDa, 79.4 kDa, and 72.4 kDa at 150 mM (brown curve), 300 mM (blue curve), 500 mM (pink curve), and 1 M NaCl (black curve), respectively. **(ii)** SDS-PAGE analysis of elute fraction corresponding to each of the GFC peaks. Lane M: Molecular weight markers; Lane 1-4: Elute fractions corresponding to the peaks from GFC at 150 mM, 300 mM, 500 mM, and 1 M NaCl, respectively. **F. Protein concentration dependent oligomerization of CaVps75Δ_C_. (i)** GFC profile for CaVps75Δ_C_ eluted from HiLoad 16/60 Superdex 200 column at 67.2 ml, 72.4 ml, and 73.1 ml which correspond to M_r_ of 117.4 kDa, 75.8 kDa, and 70.8 kDa at 50 μM (blue curve), 18 μM (pink curve), and 3.5 μM (black curve) protein, respectively. **(ii)** SDS-PAGE analysis of elute fraction corresponding to each of the GFC peaks. Lane M: Molecular weight markers; Lane 1-4: Elute fractions corresponding to the peaks from GFC at 50 μM, 18 μM, and 3.5 μM protein, respectively.

At 2 mg/ml protein concentrations, both His_6_-CaVps75 and His_6_-CaVps75Δ_C_ showed ionic strength dependent oligomerization (Figure 3B and Figure 3E, respectively). At physiological ionic strengths (150 mM NaCl), both recombinant proteins, CaVps75 and CaVps75Δ_C_ (eluted using elution buffer containing 150 mM), exhibit higher order oligomeric state, while at 300 mM NaCl the protein shifted to tetrameric and at 500 mM and higher to a dimeric form when loaded. Further, at 300 mM NaCl, we saw a protein concentration dependent oligomerization of both CaVps75 and CaVps75Δ_C_ (Figure 3 C and F, respectively). These results are consistent with what has been reported by Mingzhu Wang’s group [23] and confirm that, (i) the oligomerization is dictated at least in part by electrostatic interactions, and (ii) the acidic C-terminal domain of CaVps75 is not crucial for this interaction.

## Cloning, expression and purification of CaAsf1

His_6_-CaAsf1 was cloned in pCold I vector between XhoI and BamHI site and confirmed by plasmid PCR (Figure S1 E). His_6_-CaAsf1 was expressed in Rosetta(DE3) and purified using Ni^+2^-NTA affinity chromatography followed by GFC (Figure 4), as explained in the Methods section. The GFC HiLoad 16/60 Superdex 75 column used was calibrated using protein standards (Figure S2 B). The gel filtration profile obtained suggests that recombinant CaAsf1 purifies as a monomer under these conditions.

**Figure 4.**
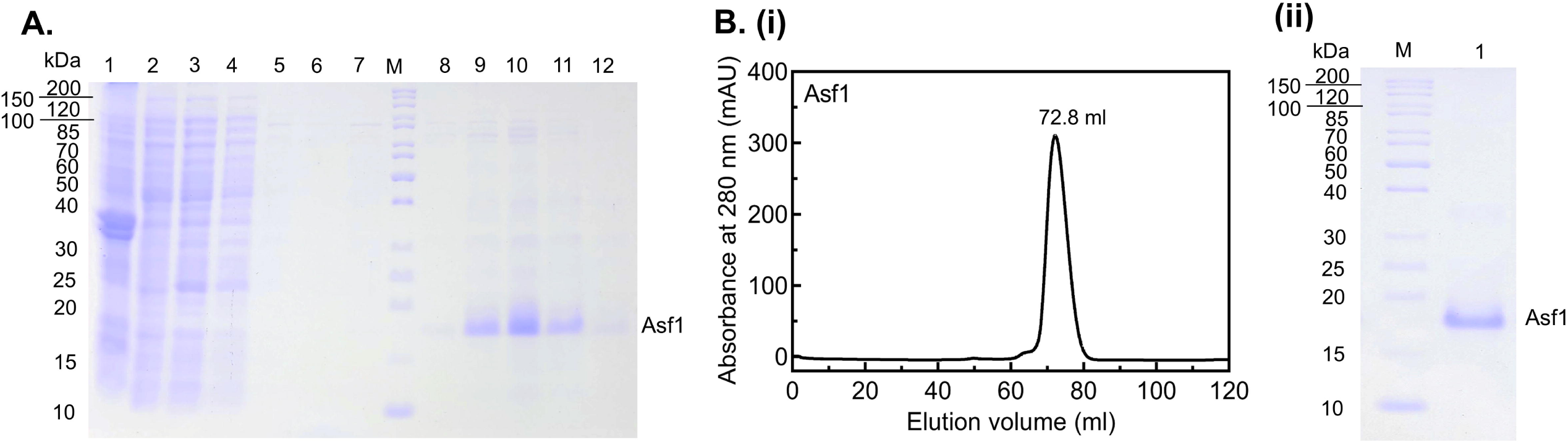
Purification of CaAsf1. *E. coli* Rosetta(DE3) cells transformed with the respective recombinant plasmid were grown until OD_600nm_ ∼0.6 and protein expression induced using 0.5 mM IPTG at 16 °C for 16 h in. The cells were harvested and lysed in 50 mM HEPES-KOH, pH 7.5, containing 150 mM NaCl, 0.5 mM DTT, 5% glycerol, and 1 mM PMSF. **A. SDS-PAGE analysis of the expression and purification of CaAsf1.** Samples from different steps of the Ni^+2^-NTA affinity purification of CaAsf1 were analyzed using 12% SDS-PAGE. Lanes 1 and 2: Pellet and supernatant fractions, respectively, obtained after centrifugation of the cell lysate; Lane 3: Flow-through fraction from the Ni^+2^-NTA column, showing the unbound proteins; Lanes 4 and 5: wash fractions with buffer containing 150 mM NaCl and 500 mM NaCl, respectively; Lane 6: wash fraction with buffer containing 150 mM NaCl with 10 mM and 20 mM imidazole, respectively; Lane M: Molecular weight markers; Lanes 8-12: protein fractions (2 ml each) eluted in the presence of 200 mM imidazole. **B. GFC analysis of purified CaAsf1.** The eluted fractions (8-12) were pooled and concentrated and loaded onto a HiLoad 16/60 Superdex 200 column. **(i)** GFC profile for CaAsf1 shows one peak at 72.8 ml volume, corresponding to the expected M_r_ of a monomer (19 kDa). **(ii)** SDS-PAGE analysis of the peak obtained at 72.8 ml in GFC. Lane M: Molecular weight markers; Lane 1: GFC peak fraction.

## Recombinant CaRtt109 forms a stable complex with CaVps75 but not with CaAsf1

To see whether CaRtt109 and CaVps75 interact, purified CaRtt109 was incubated overnight with 4-fold molar excess of CaVps75, followed by GFC analysis. Both proteins eluted in a single peak, indicating a stable protein complex formation (Figure 5A; Figure S3 A). A similar observation was made for CaRtt109 and CaVps75Δ_C_ (Figure 5B; Figure S3 B). On the other hand, CaRtt109Δ_Loop_ preincubated with either CaVps75 or CaVps75Δ_C_ eluted in separate peaks in GFC (Figure 5 C-D; Figure S3 C-D), suggesting weakened interaction between the HAT and its chaperone in the absence of the 118-160 loop region of CaRtt109.

**Figure 5.**
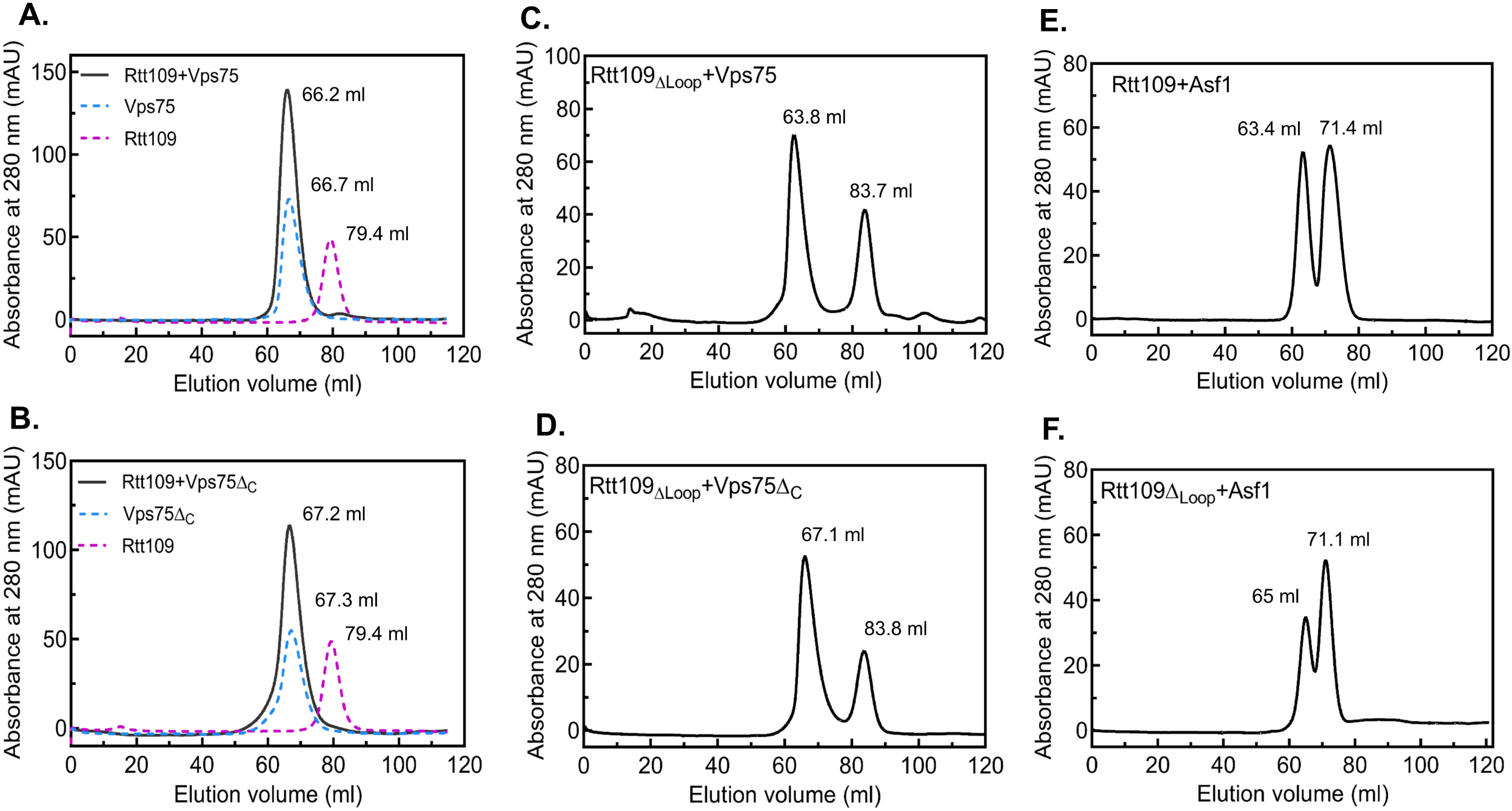
**Interaction of CaRtt109 with its chaperones. A. Co-elution of CaRtt109 with CaVps75**. Gel filtration chromatography (GFC) profile for CaRtt109+CaVps75 mixture (black curve) performed using HiLoad 16/60 Superdex 200 column showing one peak obtained at 66.2 ml elution volume, corresponding to M_r_ of 128.8 kDa. The GFC profile for CaVps75 alone (eluted at M_r_ of 123 kDa) and CaRtt109 alone (eluted at M_r_ of 41.6 kDa), depicted in blue and magenta, respectively, are also shown for comparison. **B. Co-elution of CaRtt109 with CaVps75Δ_C_.** GFC profile for CaRtt109+CaVps75Δ_C_ mixture (black curve) performed using HiLoad 16/60 Superdex 200 column showing one peak obtained at 67.2 ml elution volume, corresponding to M_r_ of 118.3 kDa. The GFC profile for CaVps75Δ_C_ alone (M_r_ of 117.5 kDa) and Rtt109 alone (M_r_ of 41.6 kDa), depicted in blue and magenta, respectively, are also shown for comparison. **C. Co-elution of CaRtt109Δ_Loop_ with CaVps75.** GFC profile for CaRtt109Δ_Loop_+CaVps75 mixture performed using HiLoad 16/60 Superdex 200 column showing two peaks obtained at 63.8 ml and 83.7 ml elution volume, corresponding to M_r_ of 154.8 kDa and 31.1 kDa, respectively. **D. Co-elution of CaRtt109Δ_Loop_ with CaVps75Δ_C_.** GFC profile for CaRtt109Δ_Loop_+CaVps75Δ_C_ mixture performed using HiLoad 16/60 Superdex 200 column showing two peaks obtained at 67.1 ml and 83.8 ml elution volume, corresponding to M_r_ of 119.4 kDa and 30.9 kDa, respectively. **E. Co-elution of CaRtt109 with CaAsf1.** GFC profile for CaRtt109+CaAsf1 mixture performed using HiLoad 16/60 Superdex 75 column showing two peaks obtained at 63.4 ml and 71.4 ml elution volume, corresponding to M_r_ of 34.3 kDa and 22 kDa, respectively. **F. Co-elution of CaRtt109Δ_Loop_ with CaAsf1.** GFC profile for CaRtt109Δ_Loop_+ CaAsf1 mixture performed using HiLoad 16/60 Superdex 75 column shows two peaks obtained at 65 ml and 71.1 ml elution volume, corresponding to M_r_ of 31.6 kDa and 22.3 kDa, respectively.

Likewise, CaRtt109 or CaRtt109Δ_Loop_ preincubated with 2-fold molar excess of CaAsf1 were loaded onto a HiLoad Superdex 75 pg GFC column for analysis. Both proteins eluted in separate peaks rather than together (Figure 5 E-F; Figure S3 E-F), suggesting weak or no interaction between CaRtt109/ CaRtt109Δ_Loop_ and CaAsf1.

The strength of these interactions was quantified by Biolayer Interferometry (BLI), which measures the real-time association and dissociation between a ligand and the analyte. Here too, CaRtt109 was found to bind strongly with both CaVps75 and CaVps75Δ_C_ (Figure 6 A-B) with dissociation constants (K_D_) of 21.8 ± 0.6 nM and 37.4 ± 0.9 nM, respectively (assuming dimeric form of CaVps75 as most the prominent one in the solution at nanomolar protein concentrations and 300 mM NaCl). In the case of CaRtt109Δ_Loop_, the strength of the interaction was reduced by ∼20 folds for CaVps75 and by ∼13 folds for CaVps75Δ_C_ (Figure 6 C-D). Thus, we conclude that the 118-160 residue loop of CaRtt109 is required for its interaction with CaVps75.

**Figure 6.**
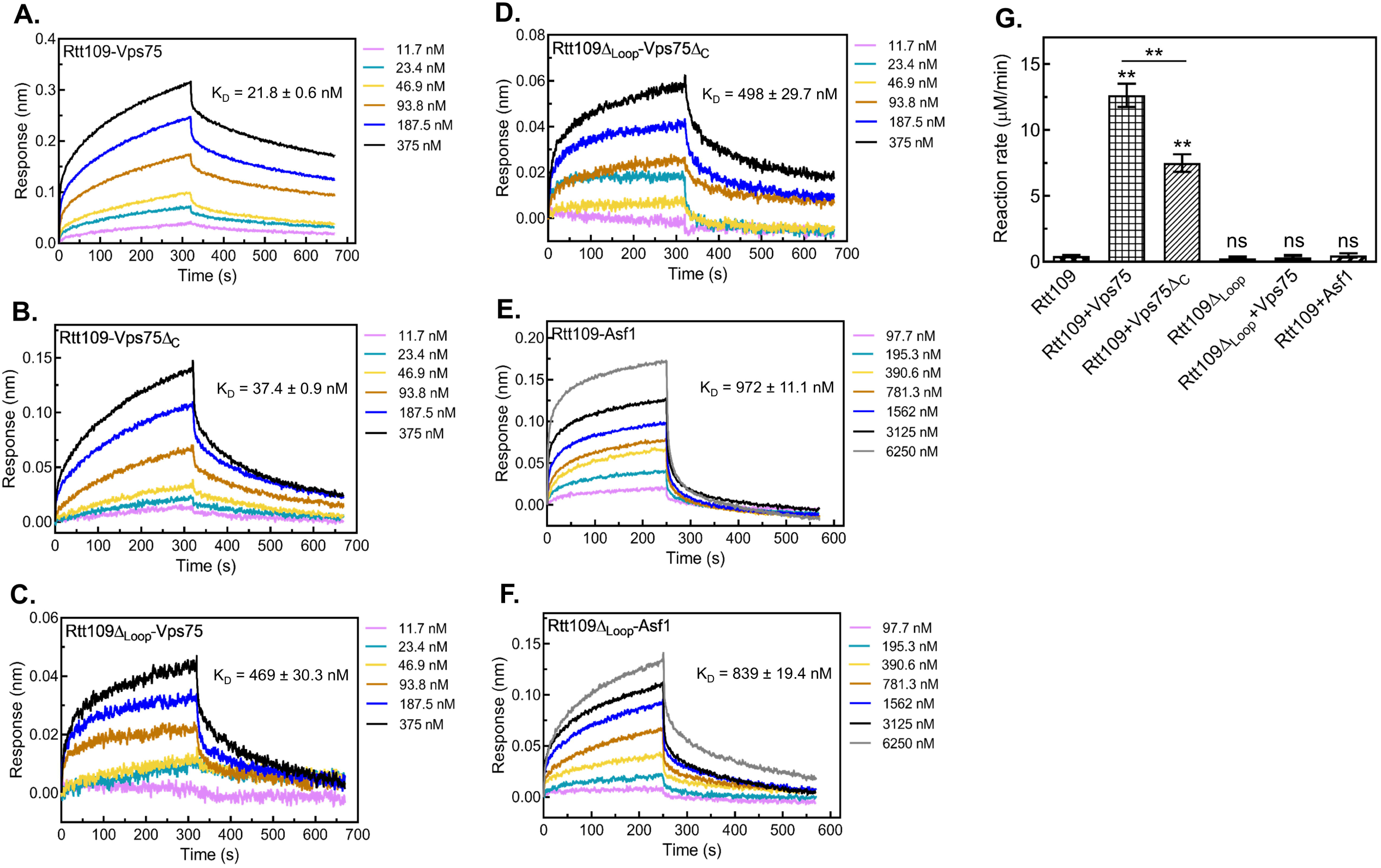
Binding analysis and catalytic activity of CaRtt109 with its chaperones. The binding of CaRtt109 with its chaperones was studied using biolayer interferometry (BLI). **A.** and **B.** shows the wavelength shift (response) upon association and dissociation of immobilized CaRtt109 to CaVps75 and CaVps75Δ_C_, respectively. **C.** and **D.** shows the wavelength shift (response) upon association and dissociation of immobilized CaRtt109Δ_Loop_ to CaVps75 and CaVps75Δ_C_, respectively. **E.** and **F.** shows the wavelength shift (response) upon association and dissociation of immobilized CaRtt109 and CaRtt109Δ_Loop_ to CaAsf1, respectively. Each concentration of analyte protein is depicted in a different color. **G. Activity of CaRtt109 towards H3 peptide monitored spectrophotometrically using a pyruvate dehydrogenase coupled assay.** The assay was performed in the presence of 200 μM H3 peptide and 50 μM AcCoA, at 0.5 µM CaRtt109 with 2 µM CaAsf1, 2 µM CaVps75, 2 µM CaVps75Δ_C_, 0.5 µM CaRtt109Δ_Loop_, where added. Activity of Rtt109 was detected only in presence of Vps75, which was significantly reduced for CaRtt109 with CaVps75Δ_C_. The average of initial reaction rates were plotted with standard deviation from three different experiments. The ‘ns’ denote ‘non-significant’, while symbol ** show significance with p-value <0.01.

Both CaRtt109 and CaRtt109Δ_Loop_ showed significantly weaker association with Asf1 as compared to CaVps75 (Figure 6 E-F). As indicated by the K_D_ values, the presence or absence of the 118-160 loop region did not greatly affect the affinity of CaRtt109 for CaAsf1, clearly showing that the interactions of CaRtt109 with its two chaperones are distinct.

## Pre-steady state kinetics of acetylation by CaRtt109 in complex with its chaperones

A coupled pyruvate dehydrogenase assay was utilized to study the activity of CaRtt109. A synthetic histone N-terminal peptide composed of 20 residues of *C. albicans* H3 was used as the substrate. Since ScGcn5 HAT domain was previously reported to be active in a similar assay using a synthetic N-terminal H3 peptide substrate H3 (for H3K14 acetylation) [24]. ScGcn5(99-262) was cloned, expressed and purified as explained in Methods and used as the positive control for the assay (Figure S4 A-C).

Next, CaRtt109 alone and in combination with its chaperones was used in the HAT assay and the initial reaction rates recorded (Figure 6G). CaRtt109 by itself showed no detectable activity, but in the presence of CaVps75 it was able to acetylate the peptide substrate. However, the CaRtt109-CaVps75Δ_C_ combination showed a drastic reduction in the reaction rates as compared to CaRtt109-CaVps75. CaRtt109Δ_Loop_, showed no activity in the presence of CaVps75, which might be a reflection of the reduced stability of the CaRtt109Δ_Loop_-CaVps75 seen above. The addition of CaAsf1 to CaRtt109 did not activate acetylation. Given that recombinant CaRtt109 did not also form a stable complex with CaAsf1, this was not surprising.

## *In silico* screening of potential inhibitor molecules

Structure-based virtual screening of the Life Chemicals compound library was performed using the Genetic Optimization for Ligand Docking (GOLD) version 5.1 software. CaRtt109 homology model was generated using crystal structure of *S. cerevisiae* Rtt109-Vps75 complex (PDB ID: 3Q68) [25] as a template and was used as the receptor model for the docking studies. Docked compounds were ranked based on their GOLD Score and X-SCORE values [26,27]. The ten highest-ranked compounds were selected by considering both their predicted docking scores and the nature of their interactions with residues within the active site of Rtt109 (Table I). Protein ligand interactions were analyzed and visualized using Discovery Studio Visualizer, in which the protein was defined as the receptor and each screened compound was defined as the ligand to generate two-dimensional interaction diagrams (Figure 7). The predicted binding modes indicated that the compounds interact with the Rtt109 active site primarily through hydrophobic interactions, hydrogen bonds, and van der Waals contacts.

**Figure 7:**
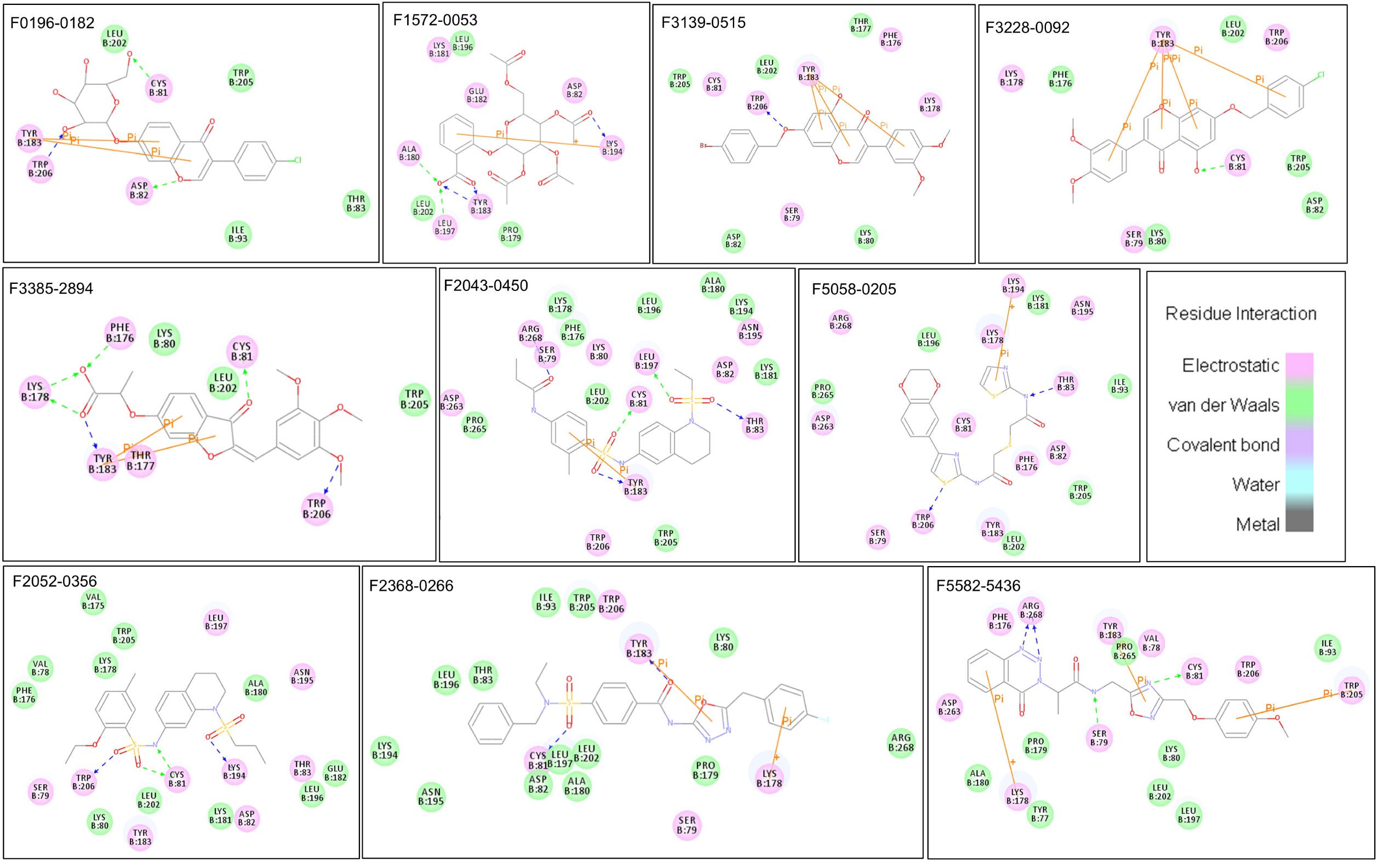
Illustration of the interactions between potential inhibitors with the active site residues of CaRtt109 upon docking using GOLD software. The docking was performed using CaRtt109 structure model [prepared using ScRtt109-ScVps75 structure (PDB ID 3Q68) as a template]. The different modes of interaction between CaRtt109 and compounds were calculated using the Discovery Studio Visualizer.

**Table I.**
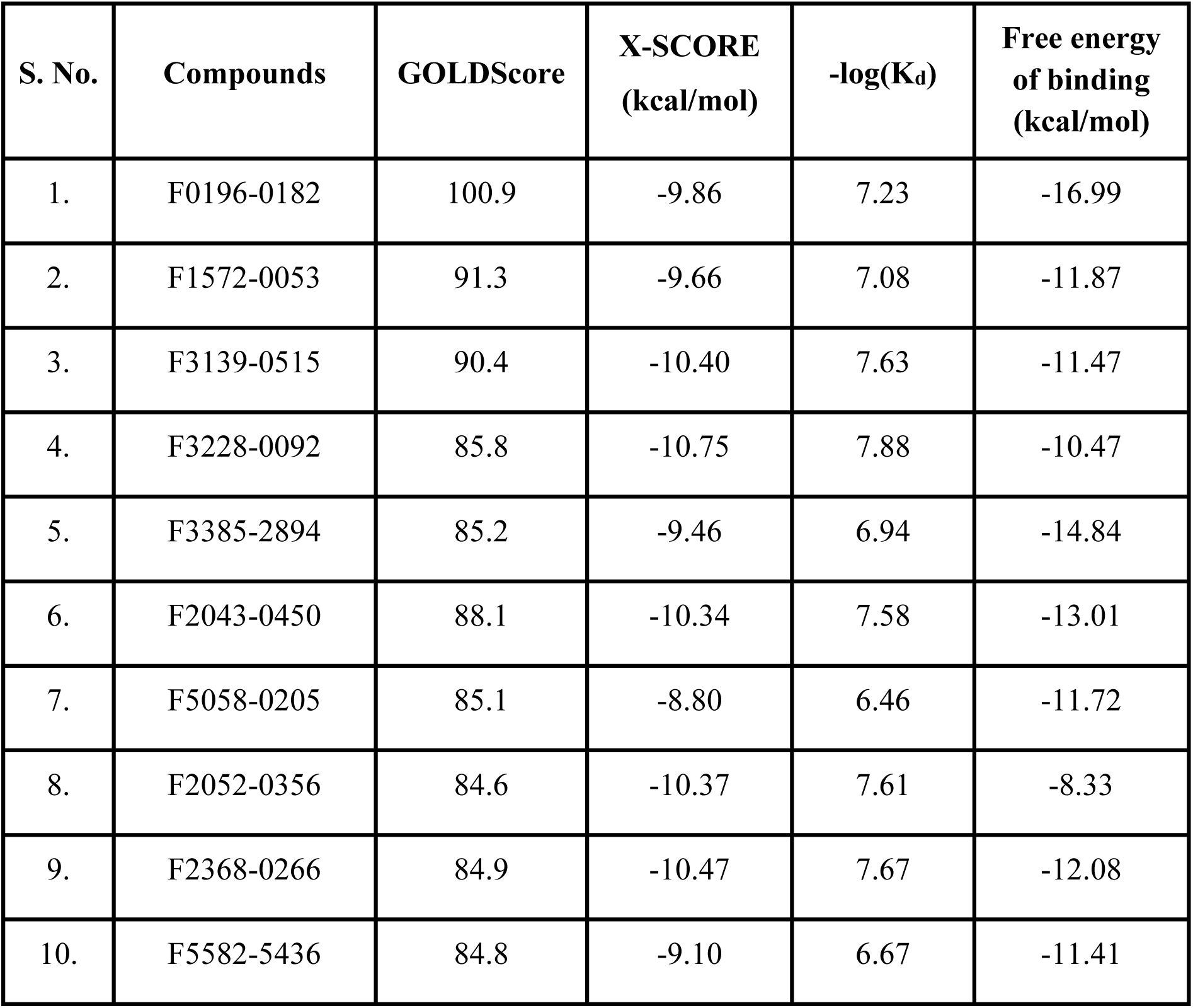
Structure-Based Docking of Life Chemicals Compounds Against the CaRtt109 Homology Model. Molecular docking was performed using the CaRtt109 homology model generated from the crystal structure of the *S. cerevisiae* Rtt109-Vps75 complex lacking acetyl coenzyme A (AcCoA) (PDB ID: 3Q68) as the structural template. Docking calculations were carried out using the Genetic Optimization for Ligand Docking (GOLD) software. During docking, the protein structure was maintained as a rigid receptor, while compounds from the Life Chemicals library served as flexible ligands. Docking was performed using the Genetic Algorithm (GA) with 10 independent docking runs per ligand and the default search algorithm parameters. Protein-ligand interactions were subsequently analyzed and visualized using Discovery Studio Visualizer. The ten highest-ranked ligands, selected based on their GOLDScore and X-SCORE values, are listed below.

| S. No. | Compounds | GOLDScore | X-SCORE<br>(kcal/mol) | -log(K<sub>d</sub>) | Free energy<br>of binding<br>(kcal/mol) |
| --- | --- | --- | --- | --- | --- |
| 1. | F0196-0182 | 100.9 | -9.86 | 7.23 | -16.99 |
| 2. | F1572-0053 | 91.3 | -9.66 | 7.08 | -11.87 |
| 3. | F3139-0515 | 90.4 | -10.40 | 7.63 | -11.47 |
| 4. | F3228-0092 | 85.8 | -10.75 | 7.88 | -10.47 |
| 5. | F3385-2894 | 85.2 | -9.46 | 6.94 | -14.84 |
| 6. | F2043-0450 | 88.1 | -10.34 | 7.58 | -13.01 |
| 7. | F5058-0205 | 85.1 | -8.80 | 6.46 | -11.72 |
| 8. | F2052-0356 | 84.6 | -10.37 | 7.61 | -8.33 |
| 9. | F2368-0266 | 84.9 | -10.47 | 7.67 | -12.08 |
| 10. | F5582-5436 | 84.8 | -9.10 | 6.67 | -11.41 |

Notably, several compounds formed interactions with highly conserved CaRtt109 residues, including Asp82, Tyr183, Trp205, and Trp206, which are known to play essential roles in acetyl coenzyme A (AcCoA) binding and catalytic activity based on studies of ScRtt109 [22,28]. In addition, interactions were observed with Phe176, Tyr183, and the aliphatic regions of Lys80 and Lys178, residues that constitute the hydrophobic tunnel proposed to function as the histone substrate-binding pocket [22,28]. These interactions suggest that the identified compounds occupy functionally important regions of the catalytic pocket and may interfere with substrate recognition or enzymatic activity. Based on commercial availability, six of the top-ranked compounds were procured for subsequent experimental evaluation.

## The compound F2368-0266 showed highest affinity for Rtt109

The interaction and binding affinity of CaRtt109 with the six shortlisted compounds were estimated in vitro using biolayer interferometry (BLI), where biotinylated CaRtt109 was immobilized onto super-streptavidin biosensors. The compounds F3139-0515, F2043-0450, F2052-0356, F3228-0092, and F3385-2894 showed lower affinity with K_D_ values in high micromolar range (207 ± 26.4 µM, 74.7 ± 4.6 µM, 109 ± 18.2 µM, 332 ± 55.4 µM, and 214 ± 16.1 µM, respectively), while F2368-0266 showed highest binding affinity with K_D_ values of 10.5 ± 1.15 µM (Figure 8; Table II). Therefore, compound F2368-0266 was tested further for its inhibitory activity in the HAT assay as described below.

**Figure 8.**
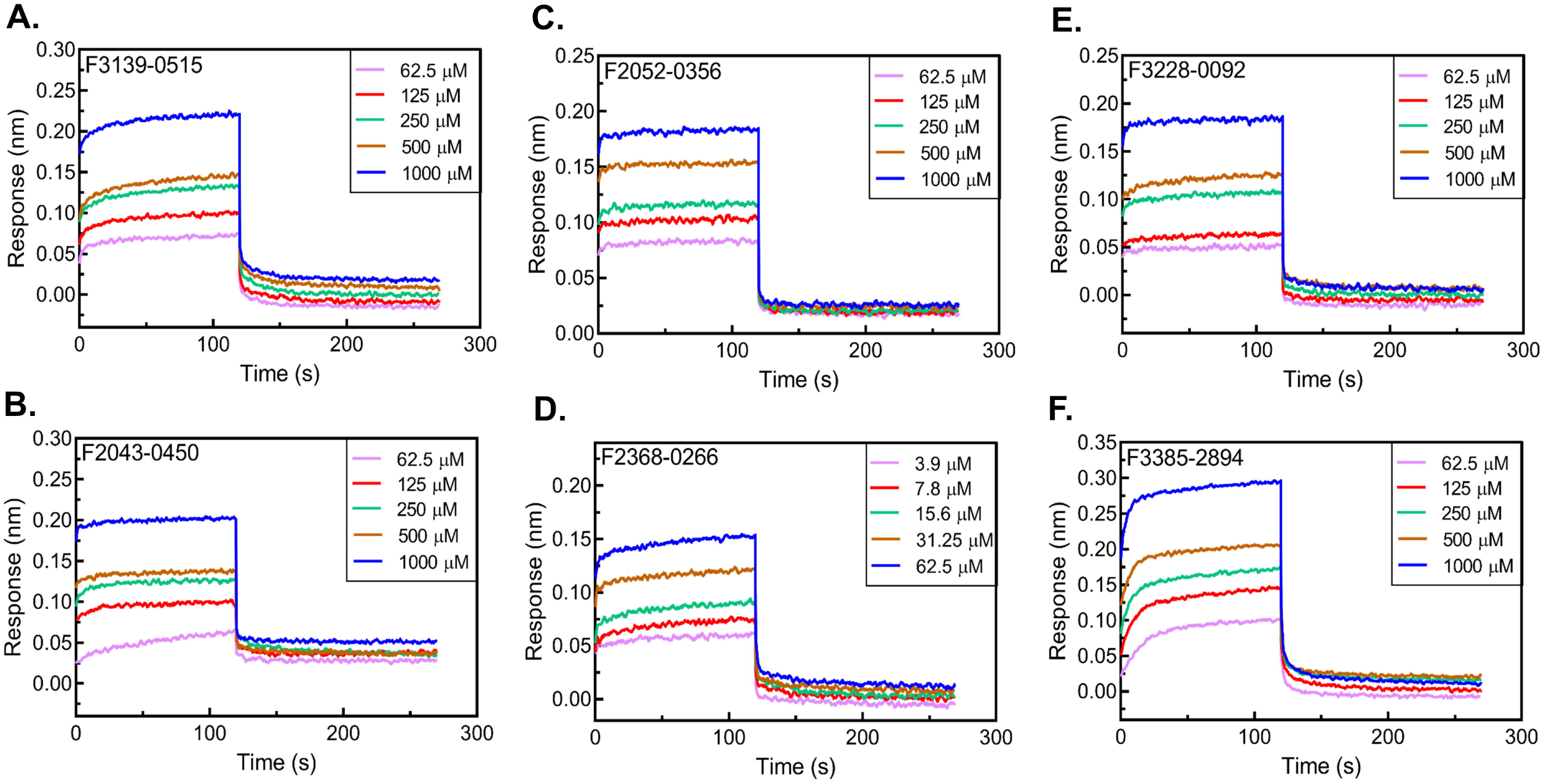
Binding study of CaRtt109 with test compounds using biolayer interferometry (BLI). The purified CaRtt109 was biotinylated and immobilized onto the super-streptavidin biosensors. The binding assay was performed thrice for each compound as analyte and the best representative graph is shown. The wavelength shift (response) upon association and dissociation of immobilized CaRtt109 to compound **A.** F3139-0515, **B.** F2043-0450, **C.** F2052-0356, **D.** F2368-0266, **E.** F3228-0092, and **F.** F3385-2894 at the indicated concentration, is shown. Each concentration of analyte protein is depicted in a different color. The control/ reference corresponding to the buffer in the absence of inhibitor was subtracted from each trace.

**Table II.** The affinity constant values (K_D_) obtained from BLI assay for CaRtt109 interaction with test compounds. The purified CaRtt109 was biotinylated and immobilized onto the super-streptavidin biosensors. The binding assay was performed thrice for each compound as analyte. The wavelength shift upon association and dissociation of immobilized CaRtt109 to each compound was recorded, and the affinity constant (K_D_) value was obtained.

| S. No. | Compound | $K_D$ (μM) |
| --- | --- | --- |
| 1. | F3139-0515 | $207 \pm 26.4$ |
| 2. | F2043-0450 | $74.7 \pm 4.6$ |
| 3. | F2052-0356 | $109 \pm 18.2$ |
| 4. | F2368-0266 | $10.5 \pm 1.15$ |
| 5. | F3228-0092 | $332 \pm 55.4$ |
| 6. | F3385-2894 | $214 \pm 16.1$ |

## Steady state kinetics of CaRtt109-CaVps75 in the absence and presence of F2368-0266

In order to study the effect of the potential inhibitor on the HAT activity of CaRtt109, steady state enzyme kinetics of the CaRtt109-CaVps75 complex was studied both in the absence and presence of F2368-0266. The assays were performed in two ways: (1) Initial reaction rates were obtained at fixed AcCoA (donor) concentration while varying histone H3 peptide (acceptor/ substrate) concentration, and (2) initial reaction rates were obtained at fixed H3 peptide concentration while varying AcCoA concentration. In this second case, at concentrations below ∼5 µM AcCoA, cloudiness was observed in the reaction, suggesting a probable requirement of AcCoA for stabilizing the H3 peptide-bound CaRtt109-CaVps75 complex in the assay conditions. Such an effect was previously reported for ScRtt109-ScVps75 in a HAT assay that used full-length H3 as substrate [19].

As before, initial rates of the reaction were obtained from the HAT assays and were used to obtain Michaelis-Menten plots as well as Lineweaver-Burk plots of the data (Figure 9 A-B) from which steady state enzyme kinetic parameters for the reaction were obtained (Table III). As can be seen from this table, the maximal reaction rates (V_max_ values) achieved by the CaRtt109-CaVps75 complex were roughly comparable irrespective of whether the concentration of the acceptor was varied or that of the donor. Further, the values of the Michaelis-Menten constant (K_M_) obtained for H3 peptide in the case of CaRtt109-CaVps75 is about 1.5 fold higher than that reported for recombinant ScRtt109-ScVps75 by Albaugh et al., 2010, and the K_M_ for AcCoA is ∼ 60-70 fold higher. In other words, the recombinant *C. albicans* HAT complex appears to have a weaker affinity for the acceptor and donor substrates as compared to its *S. cerevisiae* counterpart.

**Figure 9.**
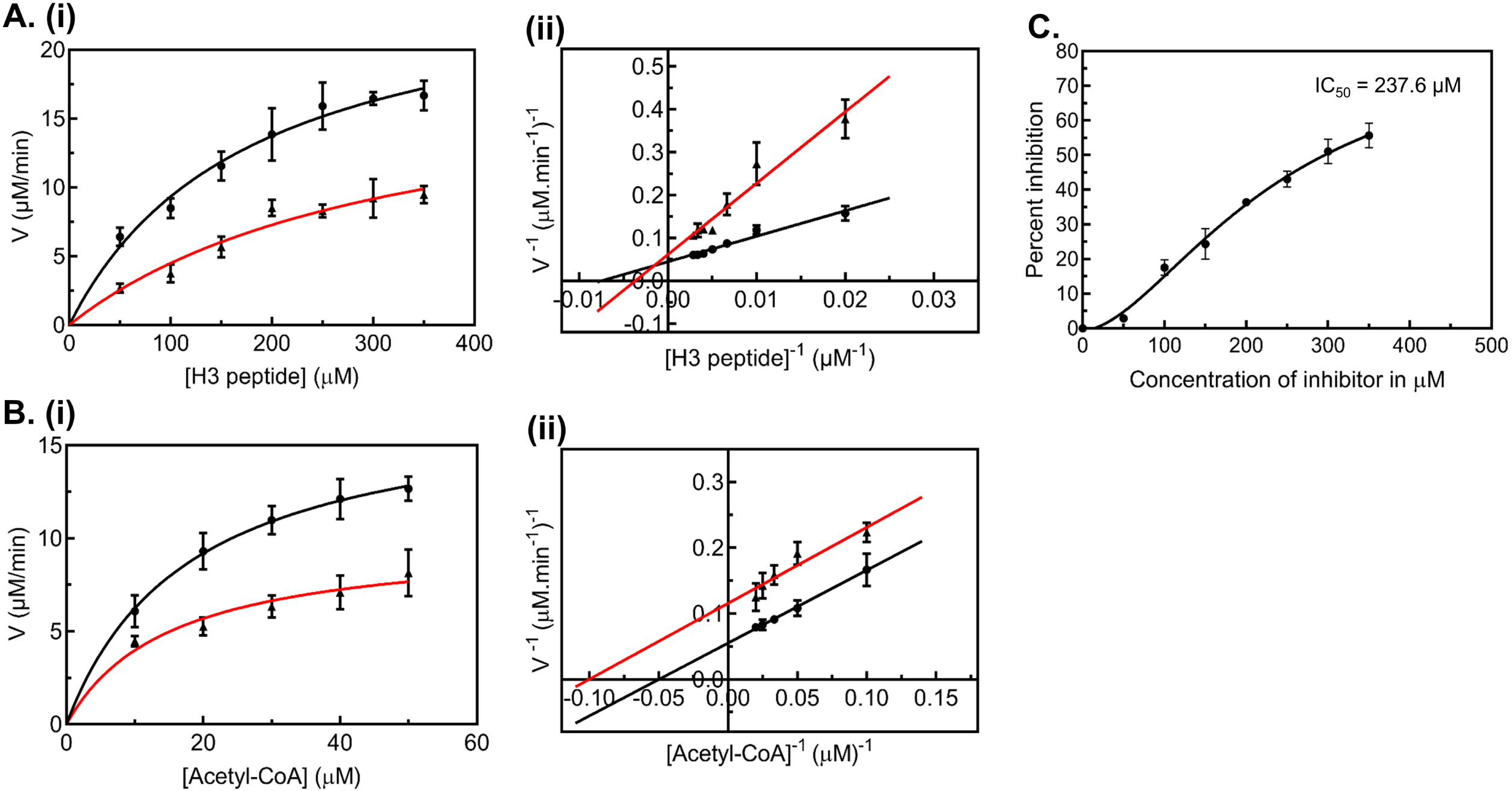
Activity of CaRtt109 and the effect of compound F2368-0266 on activity. Activity of CaRtt109 towards H3 peptide monitored spectrophotometrically using a pyruvate dehydrogenase coupled assay. The enzyme kinetics were performed with 0.5 µM CaRtt109 and 2 µM CaVps75 mixture without compound F2368-0266 (shown in black) and in presence of 300 μM inhibitor (shown in red). The reactions were performed in presence of 2% DMSO and 2% glycerol concentrations. **A.** Steady state kinetics of CaRtt109-CaVps75 catalyzed acetylation on H3 peptide at varying H3 peptide concentrations (50 to 350 μM) keeping AcCoA concentration fixed at 50 μM. The average initial reaction rates were plotted with standard deviation from three different experiments to obtain **(i)** Michaelis-Menten plot and **(ii)** Lineweaver-Burk plot. **B.** Steady state kinetics of CaRtt109-CaVps75 catalyzed acetylation on H3 peptide at varying AcCoA concentration (10 to 50 μM) keeping H3 peptide fixed at 200 μM. The average initial reaction rates were plotted with standard deviation from three different experiments to obtain **(i)** Michaelis-Menten and **(ii)** Lineweaver-Burk plots. **C.** Percent inhibition of HAT activity by F2368-0266 was plotted as a function of the inhibitor concentration (50 to 350 μM) at 200 μM H3 peptide and 50 μM AcCoA.

**Table III.**
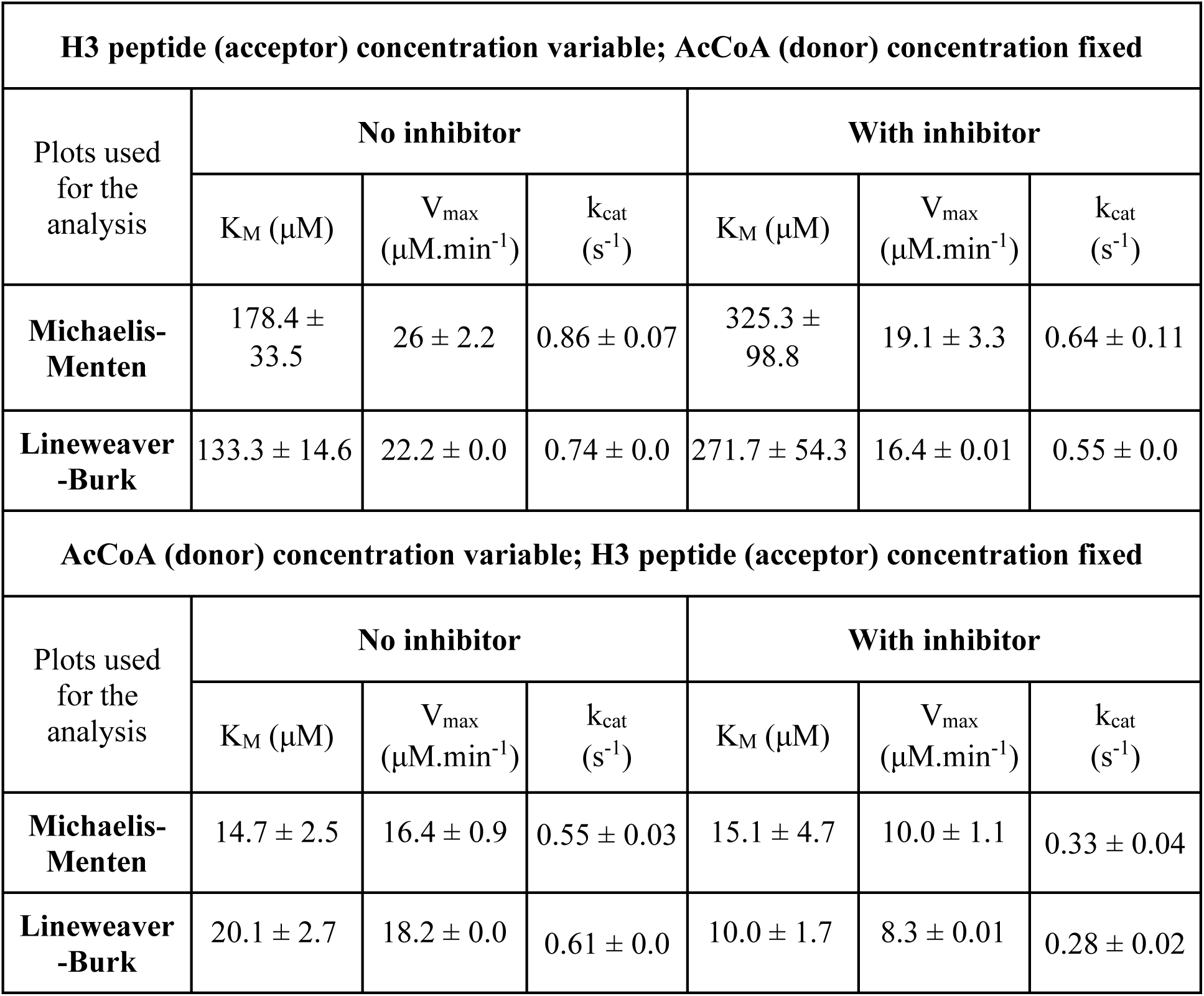
The steady state enzyme kinetic parameters obtained for HAT activity of CaRtt109-CaVps75 without and with inhibitor (F2368-0266). HAT activity was monitored spectrophotometrically using a pyruvate dehydrogenase coupled assay by mixing CaRtt109 (0.5 µM) with CaVps75 (2 µM) in the absence and presence of inhibitor F2368-0266. The reaction kinetics was monitored at varying H3 peptide concentrations (0 to 350 μM) while keeping acetyl-CoA concentration fixed at 50 μM, or by varying acetyl-CoA concentration (0 to 50 μM) and keeping H3 peptide fixed at 200 μM. The average initial reaction rates from three independent experiments were used to obtain Michaelis-Menten and Lineweaver-Burk plots. The estimated steady state kinetic parameters along with their standard deviations from these plots are listed below.

| **H3 peptide (acceptor) concentration variable; AcCoA (donor) concentration fixed** |  |  |  |  |  |  |
| --- | --- | --- | --- | --- | --- | --- |
| Plots used for the analysis | No inhibitor |  |  | With inhibitor |  |  |
| | $K_M$ ($\mu\text{M}$) | $V_{\max}$ ($\mu\text{M}.\text{min}^{-1}$) | $k_{\text{cat}}$ ($\text{s}^{-1}$) | $K_M$ ($\mu\text{M}$) | $V_{\max}$ ($\mu\text{M}.\text{min}^{-1}$) | $k_{\text{cat}}$ ($\text{s}^{-1}$) |
| **Michaelis-Menten** | $178.4 \pm 33.5$ | $26 \pm 2.2$ | $0.86 \pm 0.07$ | $325.3 \pm 98.8$ | $19.1 \pm 3.3$ | $0.64 \pm 0.11$ |
| **Lineweaver-Burk** | $133.3 \pm 14.6$ | $22.2 \pm 0.0$ | $0.74 \pm 0.0$ | $271.7 \pm 54.3$ | $16.4 \pm 0.01$ | $0.55 \pm 0.0$ |
| **AcCoA (donor) concentration variable; H3 peptide (acceptor) concentration fixed** |  |  |  |  |  |  |
| Plots used for the analysis | No inhibitor |  |  | With inhibitor |  |  |
| | $K_M$ ($\mu\text{M}$) | $V_{\max}$ ($\mu\text{M}.\text{min}^{-1}$) | $k_{\text{cat}}$ ($\text{s}^{-1}$) | $K_M$ ($\mu\text{M}$) | $V_{\max}$ ($\mu\text{M}.\text{min}^{-1}$) | $k_{\text{cat}}$ ($\text{s}^{-1}$) |
| **Michaelis-Menten** | $14.7 \pm 2.5$ | $16.4 \pm 0.9$ | $0.55 \pm 0.03$ | $15.1 \pm 4.7$ | $10.0 \pm 1.1$ | $0.33 \pm 0.04$ |
| **Lineweaver-Burk** | $20.1 \pm 2.7$ | $18.2 \pm 0.0$ | $0.61 \pm 0.0$ | $10.0 \pm 1.7$ | $8.3 \pm 0.01$ | $0.28 \pm 0.02$ |

When the assay was carried out in the presence of F2368-0266 we noted that the inhibitor reduced the affinity (increased K_M_) of CaRtt109-CaVps75 complex for the H3 peptide substrate by roughly two-fold while not drastically altering the V_max_ values (Figure 9 A-B; Table III). As a result, the turnover number (k_cat_) too was reduced. On the other hand, it did not appear to affect the binding of AcCoA to the protein complex. In other words, the inhibitor appears to specifically and competitively inhibit substrate binding.

The calculated concentration at which the HAT activity of CaRtt109-CaVps75 complex is inhibited up to 50% (IC_50_) by F2368-0266 is 237.6 μM (Figure 9C). Application of the Cheng-Prusoff equation, K_i_ = IC_50_/(1+ [I]/K_D_ [29], under these assumptions yields an inhibition constant (K_i_) of ∼ 112 μM for F2368-0266, which implies only a slightly better affinity for the inhibitor as compared to the peptide substrate.

It is also worth noting that the affinity of CaRtt109-CaVps75 for AcCoA is over 10-fold better (lower K_M_ values) than for the H3 peptide. In other words, if both AcCoA and the H3 peptide are present in the reaction assay at equal concentrations, it is quite likely that the former would bind to the enzyme complex before the latter. This is also likely to be the case when both F2368-0266 and AcCoA are present simultaneously.

Hence, rather than using the ScRtt109-ScVps75 crystal structure obtained in the absence of AcCoA (3Q68) [25] which was used for the screening of inhibitors, we chose one that was obtained in the presence of AcCoA (3Q35), and generated a homology model for CaRtt109 from this. The AcCoA was placed in the AcCoA binding pocket in the CaRtt109 model and energy was minimized. Thereafter, docking the inhibitor to this structure yielded a binding site for the inhibitor that was proximal to but distinct from that for AcCoA (Figure 10). Several residues of CaRtt109 interacted with the inhibitor, including Val78, Ser79, Phe176, Lys181, Glu182, Tyr183, Ala189, Asp263, Asp264, and Pro 265, all of which are part of the hydrophobic tunnel previously proposed to interact with the histone substrate in ScRtt109 to correctly position the Lys residue for acetylation [22,28].

**Figure 10.**
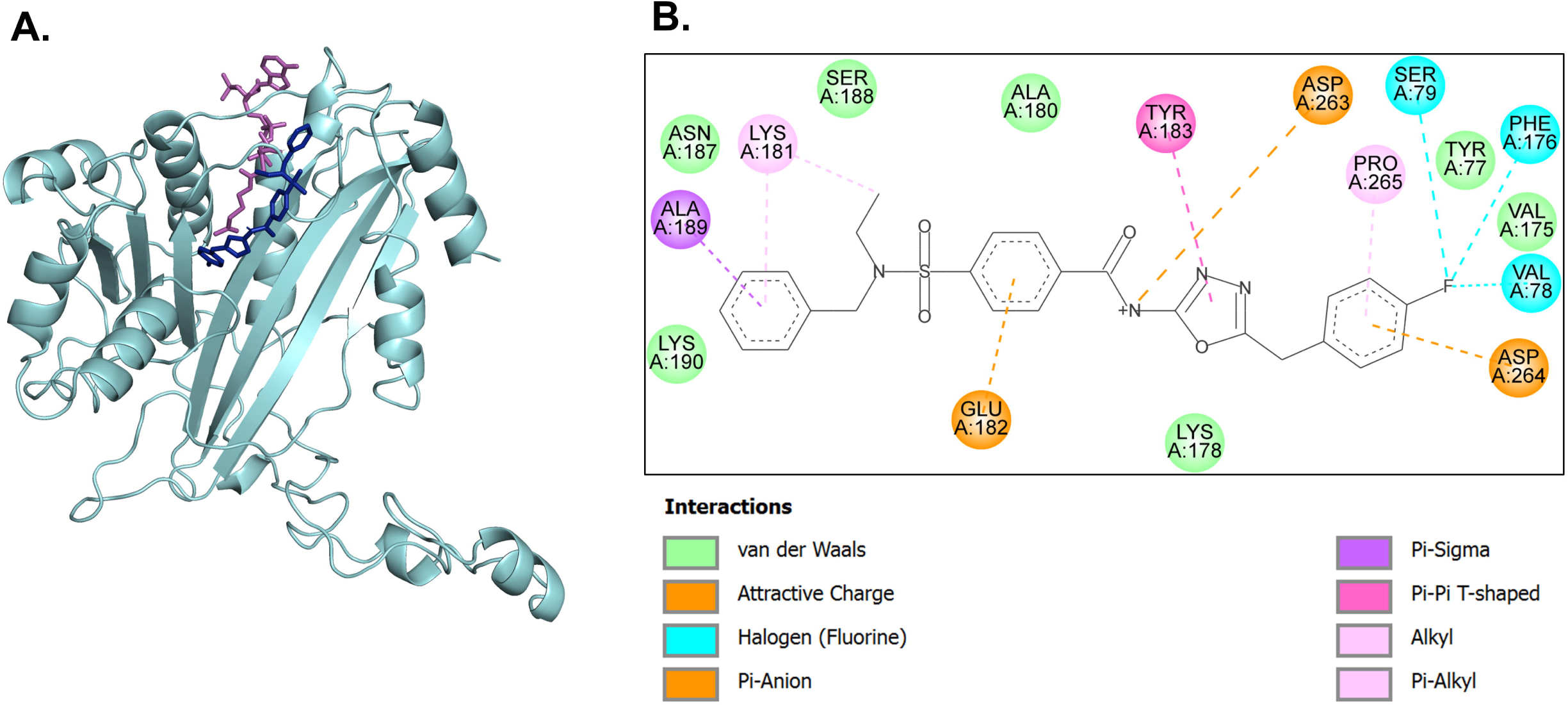
Docking of F2368-0266 in the homology model of CaRtt109 with AcCoA using AutoDock Tools 4.2. The CaRtt109 homology model was generated using the crystal structure of the ScRtt109-ScVps75 complex bound to AcCoA (PDB:3Q35) as the template. AcCoA was placed in the AcCoA binding pocket in the model and energy was minimized. The compound was then docked in this model. **A.** The compound F2368-0266 (shown in blue) docked proximal to the AcCoA moiety (shown in magenta) with a △G = -7.06 kcal/mol. **B.** The F2368-0266 binding pocket of CaRtt109 along with the interacting residues and the nature of the interaction are shown.

## Discussion

Epigenetic regulation of chromatin dynamics via histone modifications is one of the major mechanisms by which eukaryotic cells control gene expression, DNA repair, and maintenance of genome stability. Of the different covalent modifications possible on histone proteins, acetylation carried out by HATs, and particularly H3K56 acetylation, is crucial for controlling gene expression, morphogenesis, virulence and regulating drug response in fungi. That Rtt109 is the only HAT capable of effecting this modification in fungal cells, and that it is exclusive to the fungal kingdom, makes it a rather unique protein to study even from an antifungal drug target point-of-view.

Such a case can be specifically made out for Rtt109 from *C. albicans*, a commensal fungus capable of opportunistic infections under immunocompromised conditions and responsible for a major fraction of invasive candidiasis cases globally. Loss of Rtt109 is not lethal for *C. albicans* but it does severely attenuate its drug susceptibility as well as pathogenesis [14,30]. Further, since it regulates the first step of GPI biosynthesis, it reduces the expression of GPI anchored proteins and virulence factors at the cell surface, and alters sterol biosynthesis as well as cell wall integrity, all of which also contribute to the reduced virulence [15,16]. A large fraction of antifungals currently in use target sterol or cell wall biogenesis or directly disrupt cell membrane and cell wall integrity; but drug resistance has been rampant [31]. Newer drugs and targets, including fosmanogepix aimed at the GPI biosynthetic pathway are currently under development for precisely this reason [32,33]. Given its central role regulating all of these pathways simultaneously, *C. albicans* Rtt109 could also prove to be a good target. With this context in mind, the HAT activity of recombinant CaRtt109 was characterized in this study.

The catalytic activity of Rtt109 requires the protein to be bound to at least one of its two chaperones, Vps75 or Asf1, in *S. cerevisiae*. Hence, for this study, CaRtt109, CaVps75 and CaAsf1 were separately expressed and purified from a bacterial host. Both CaRtt109 and CaAsf1 eluted as monomeric proteins when N-terminally tagged with a 6X-His and affinity-purified using a Ni^+2^-NTA column. Since ScVps75 was previously shown to exist in different oligomeric states depending on protein concentration and ionic strengths of the buffer used, we also characterized the oligomeric state of CaVps75. This enabled us to also confirm that the protein used for the interaction studies as well as the HAT activity assays was predominantly in its dimeric state. Gel filtration chromatography suggested that each monomer of CaRtt109 forms a stable complex with a Vps75 dimer under the same conditions. BLI studies indicated a high affinity interaction between the two. Deleting the acidic C-terminal tail of CaVps75 did not affect the interaction. However, deleting a flexible 118-160 loop region of CaRtt109 significantly weakened the complex formation between the two proteins, as evidenced by gel filtration studies as well as BLI. Further, unlike the CaRtt109-CaVps75 or CaRtt109-CaVps75Δ_C_ complexes, CaRtt109Δ_Loop_–CaVps75 was poorly active in the HAT assay. Thus, although not conserved in all fungal organisms, the loop region of CaRtt109 appears to be required for its interaction with its chaperone, CaVps75, and its acetylation activity. This is reminiscent of what is observed in the case of the ScRtt109-ScVps75 complex as well [11,20,21]. In those fungal species whose Rtt109 homologs lack this loop insertion, a direct interaction with Vps75 homologs also appears to be lost [34,35].

On the other hand, recombinant CaRtt109 did not seem to form a stable complex with CaAsf1, and both proteins eluted as separate peaks in GFC. BLI studies confirmed that the affinity of CaRtt109 for CaAsf1 was ∼46 fold weaker as compared to its affinity for CaVps75. Also, the presence of CaAsf1 could not activate H3 peptide acetylation by CaRtt109 in vitro. It is known that Asf1 binds to H3 and activates acetylation of H3K56 by Rtt109 only in the presence of H4 [36]. This might explain why CaAsf1-CaRtt109 is unable to use the short H3 peptide as a substrate in vitro.

*In silico* screening of biologically active compounds from the Life Chemicals library identified several potential inhibitors of *Candida albicans* Rtt109 (CaRtt109). The compounds were ranked based on their GOLD Score and X-SCORE values, and the ten highest-scoring compounds were selected as the best potential interactors. Of these, six compounds, chosen based on their commercial availability, were experimentally evaluated using biolayer interferometry (BLI). All six compounds were confirmed to bind CaRtt109 with moderate binding affinities, exhibiting dissociation constant (K_D_) values in the micromolar range. Among the tested compounds, F2368-0266, which displayed the strongest interaction with CaRtt109, was subsequently evaluated for its ability to inhibit the histone acetyltransferase (HAT) activity of the enzyme.

Steady state kinetic parameters suggested that the inhibitor did not alter the affinity of CaRtt109-CaVps75 for AcCoA and selectively inhibited the H3 peptide binding pocket. From the estimates of K_i_ (∼ 112 μM), it appears that the affinity of the CaRtt109-CaVps75 complex for the inhibitor is only slightly better than that for the peptide substrate. Nevertheless, the inhibitor allows us to define the likely peptide binding pocket. Future experiments involving crystallization of the complex with the inhibitor and its derivatives, would be useful to design improved Rtt109 inhibitors for *Candida albicans*.

Given the importance of *Candida albicans* Rtt109 in regulating virulence attributes such as hyphal morphogenesis and GPI biosynthesis, characteristics that are unique to this human pathogen, these results have significant clinical implications, particularly in a scenario where drug targets are limited and antifungal resistance is rapidly growing.

## Materials and Methods

### Reagents

The growth media components were purchased from Hi-Media (Maharashtra, India). All the primers were synthesized by either GCC Biotech (Kolkata, West Bengal, India) or Merck (Sigma-Aldrich Chemicals). DNA polymerase, restriction enzymes and T4 DNA ligase were purchased from Thermo-Fisher Scientific, plasmid DNA purification kit from Macherey-Nagel (Düren, Germany), and gel extraction kit from Qiagen (USA). The compounds used in inhibitor screening were purchased from Life Chemicals (Ontario, Canada). The peptide substrate for the HAT assay was custom synthesized by GL Biochem, Biotech Desk (Hyderabad, India). All other chemicals were procured from Sisco Research Laboratories (SRL).

### Strains and vectors

The *E.coli* DH5α strain from Merck (BGPI, India) was used for cloning, and *E.coli* Rosetta(DE3) strain from Merck (USA) was used for protein expression. The TA cloning vector (pTZ57R/T) was purchased from MBI Fermentas (USA), the pCold I expression vector from Addgene (USA) and pRSFDuet-1 expression vector from Novagen.

## Cloning of the genes of interest

The genomic DNA of BWP17 strain of *C. albicans* was used to amplify *CaRTT109*, *CaVPS75*, and *CaASF1* genes. The full-length *CaRTT109* gene was cloned in the TA cloning vector and then subcloned into the pCold I vector. *CaRTT109*Δ_Loop_ construct, in which sequences encoding residues 118-160 corresponding to a flexible loop of the full-length protein were deleted, was subcloned in pCold I vector using overlap extension PCR. *CaVPS75* and *CaVPS75*Δ_C_ (C-terminal truncated mutant encoding residues 1-227) were cloned in the MCS-1 of pRSFDuet-1 vector. *CaASF1* (encoding full length, 1-154, amino acids) was cloned in the pCold I vector.

### Cloning, expression and purification of Rtt109 and its chaperones, CaVps75 and CaAsf1

For protein expression, *E. coli* Rosetta(DE3) competent cells were transformed with the respective recombinant plasmids. The transformed cells were induced at OD_600nm_ 0.6 for protein overexpression using 0.5 mM IPTG at 16 °C and allowed to grow for 16 h. The cells were harvested at 7000 rpm for 6 min and resuspended in a buffer containing 50 mM HEPES-KOH pH 7.5, 150 mM or 300 mM NaCl, 0.5 mM dithiothreitol (DTT), 5% glycerol, and 1 mM PMSF. For CaRtt109 and CaVps75 protein constructs, 300 mM NaCl was used in buffer throughout the purification process, and for CaAsf1 purification, 150 mM NaCl was used. Cell lysis was performed by freeze-thaw method where the cell suspension was flash-frozen in liquid nitrogen and thawed at 37 °C in a water bath. Further lysis was performed by sonication (30 s ON; 45 s OFF; 6 cycles) with 25% amplitude pulses. The cell lysate was centrifuged at 17,000 rpm for 30 min. The supernatant was then passed through Ni^+2^-NTA column, pre-equilibrated with the equilibration buffer (50 mM HEPES-KOH, pH 7.5, with 300 mM or 150 mM NaCl, 0.5 mM DTT, 5% glycerol, 2 mM ATP). The elution buffer was 50 mM HEPES-KOH, pH 7.5, containing 0.5 mM DTT, 5% glycerol, 200 mM imidazole and 300 mM or 150 mM NaCl was used to elute the bound proteins. While studying the effect of ionic strength on CaVsp75 oligomerization status, the elution buffer contained either 150 mM or 300 mM NaCl, as required. For higher ionic strengths, the NaCl concentration was adjusted as required by buffer exchange after elution.

The eluted samples were analyzed on 12% SDS-PAGE. For further purification, gel filtration chromatography (GFC) was performed for which the protein was concentrated to 2 ml using Amicon centrifugal filters and centrifuged at 12,000 rpm to remove insoluble aggregates. The protein was then injected onto a gel filtration column (HiLoad 16/60 Superdex 75 pg or 200 pg column, specified for each protein in the results section). The peak fractions were analyzed by 12% SDS-PAGE.

### Cloning, expression and purification of *S. cerevisiae* Gcn5 HAT domain

The genomic DNA of *S. cerevisiae* strain YPH500 was used to amplify *GCN5* gene sequence encoding the HAT domain (99-262 amino acid residues). The gene segment was cloned between XhoI and BamHI restriction sites of the pCold I vector. Positive clones were confirmed by double restriction digestion. The protein was over-expressed with an N-terminal His_6_-tag and purified using Ni^+2^-NTA affinity chromatography, following the same method as described in the above section. The protein was eluted in 50 mM HEPES-KOH (pH 7.5) buffer containing 150 mM NaCl, 0.5 mM DTT, and 5% glycerol. The eluted fractions were analyzed on 12% SDS-PAGE.

## Gel filtration for protein-protein interaction studies

The proteins were purified separately as described in the above section using Ni^+2^-NTA affinity chromatography followed by GFC. To study the interaction of CaRtt109 with CaVps75, 10 µM Rtt109 was incubated with 40 µM Vps75 at 4 °C overnight in 50 mM HEPES-KOH (pH 7.5) buffer containing 300 mM NaCl, 0.5 mM DTT, and 5% glycerol. The mixture was centrifuged at 12,000 rpm to remove any insoluble aggregates. It was then loaded onto a HiLoad 16/60 Superdex 200 pg column. The GFC profile was recorded and peak fractions were analyzed on SDS-PAGE. Similar steps were followed to study interaction of CaRtt109 with CaVps75Δ_C_, and CaRtt109Δ_Loop_ with both CaVps75 and CaVps75Δ_C_.

To study the interaction of CaRtt109 with CaAsf1, CaRtt109 was subjected to buffer exchange (50 mM HEPES-KOH, pH 7.5, containing 150 mM NaCl, 0.5 mM DTT, and 5% glycerol) using Amicon centrifugal filters. Thereafter, 10 µM CaRtt109 was incubated with 20 µM CaAsf1 at 4 °C overnight. The mixture was centrifuged at 12,000 rpm to remove any insoluble aggregates and loaded onto the HiLoad 16/60 superdex 75 pg column. The GFC profile was recorded and peak fractions were analyzed on SDS-PAGE. Likewise, the interaction of CaRtt109Δ_Loop_ with CaAsf1 was studied.

## Biolayer interferometry (BLI) for protein-protein and protein-ligand interactions

The biolayer interferometry experiments were conducted on the Sartorius Octet R2 system. The purified CaRtt109 protein was biotinylated using EZ-Link NHS-LC-Biotin (Thermo Scientific). The biotinylated protein (ligand) was immobilised on two streptavidin biosensors (SA) pre-hydrated in the assay buffer (the test and reference biosensor). Any free streptavidin binding sites were blocked using a 10 µg/ml biocytin solution in the assay buffer. The assay buffer composition used for studying CaRtt109 interaction with its chaperones was 20 mM HEPES-KOH pH 7.5, 2 mM β-mercaptoethanol, and 2% glycerol, containing 200 mM NaCl for CaAsf1 and 300 mM for CaVps75. For BLI studies involving small molecules as analyte, CaRtt109 was immobilized on super-streptavidin biosensors (SSA) and the assay buffer used contained 20 mM HEPES-KOH pH 7.5, 2 mM β-mercaptoethanol, 2% glycerol, and 5% DMSO. The inhibitor stock solutions were prepared in 100% DMSO and further dilutions were made in the assay buffer such that the final DMSO concentration did not exceed 5%.

The binding experiment had three steps: establishing baseline in assay buffer, an association step where the test biosensor was dipped in analyte solution, and a dissociation step in assay buffer. The experiment was performed at 20-25 °C. As a negative control, the assay buffer without analyte was used for both association and dissociation steps to normalize the data. Octet BLI discovery 12.2 software was used for data acquisition and Octet analysis 12.1 software was used for data analysis (global fitting, 1:1 binding model). All experiments were performed thrice. The binding trend remained consistent and the best representative graphs are shown in the figures. The dissociation constant (K_D_) is calculated from the ratio of the rate constants for the association (k_a_) and dissociation (k_d_) steps.

## Histone acetyltransferase (HAT) assay

A pyruvate dehydrogenase coupled spectrophotometry-based assay is utilized to study the HAT activity of CaRtt109 alone and in complex with its chaperones. The assay used is a modified version of previously reported protocols [19,24,37]. Acetyl-Coenzyme A (AcCoA) was used as the acetyl donor, and CoA generated from the HAT reaction was used as the substrate for a coupled pyruvate dehydrogenase (PDH) catalyzed reaction to produce NADH which was monitored spectrophotometrically at 340 nm [38]. The peptide substrate to be acetylated in the HAT assay, corresponding to 20 N-terminal amino acid residues of *C. albicans* histone H3 (ARTKQTARKSTGGKAPRKQL), was custom synthesized (purity >98%). A 300 µl reaction mixture contained 50 mM HEPES-KOH pH 7.5, 1 mM DTT, 2% final DMSO, 2% glycerol, 5 mM MgCl_2_, 2.4 mM TPP, 0.2 mM NAD^+^, 5 mM pyruvate, and 0.3 mg/ml PDH, AcCoA and 0.5 µM Rtt109 with 2 µM Vps75. It was incubated for 5 min at 25 °C, after which the reaction was initiated by adding histone H3 peptide and monitored for 5 min. The initial reaction rate was calculated for the linear part of the reaction by dividing absorbance change over time with NADH extinction coefficient (ɛ = 6.23 mM^-1^ cm^-1^) at 340 nm. His_6_-ScGcn5(99-262) was used as a positive control for the HAT assay.

To study the acetylation activity in the presence of the inhibitor, 300 µM of F2368-0266 was also added to the reaction mixture such that the DMSO concentration did not exceed 2%. The absorbance at 340 nm was recorded continuously every 6 s in BioTek Cytation 5 imaging reader for 5-10 min at 25 °C. To estimate the background reaction rate, a control reaction without H3 peptide was set whenever a reaction component was varied. The background reaction was then subtracted from the actual enzyme reaction where all components were present to obtain the rate of enzyme catalyzed reaction. The initial rates were calculated for reactions (with and without F2368-0266) with H3 peptide concentration varied from 50 to 350 μM at fixed AcCoA concentration of 50 μM, and with fixed H3 peptide concentration of 200 μM while varying AcCoA concentration from 10 to 50 μM. The reactions were performed thrice, each set in duplicate. The data was fit to a Michaelis-Menten plot using GraphPad Prism 8.0.2 software and steady state kinetic parameters were calculated.

## *In silico* screening of potential ligands/ inhibitors

A virtual screening was performed using approximately 1.35 million commercially available small molecules from the Life Chemicals database. The three-dimensional structure of *Candida albicans* Rtt109 (CaRtt109) was generated by homology modeling using the *S. cerevisiae* Rtt109–Vps75 complex (PDB ID: 3Q68) [25] as the template in MODELLER (https://salilab.org/modeller/) [39]. The modeled structure was prepared by adding polar hydrogen atoms and subjected to energy minimization using Swiss-PDB Viewer (https://spdbv.unil.ch/) with steepest descent and conjugate gradient algorithms to eliminate steric clashes and optimize structural geometry. Model quality and stereochemical integrity were evaluated using PROCHECK [40].

The compound library was prepared using the LigPrep module of Schrödinger to generate energy-minimized three-dimensional conformations with appropriate protonation states and stereochemistry. Molecular docking was performed using GOLD v5.1 [26], treating the protein as a rigid receptor while allowing full ligand flexibility through the Genetic Algorithm, with ten independent docking runs conducted for each ligand using default search parameters [41,42]. Protein-ligand interactions were analyzed using Discovery Studio Visualizer (https://www.3ds.com/products/biovia/discovery-studio/visualization), and compounds were ranked based on GOLD Score and X-Score (https://ics.uci.edu/∼dock/manuals/xscore1.1_manual/intro.html) binding affinity predictions. The highest-ranked compounds were prioritized based on consensus docking scores, interactions with catalytic-site residues, and chemical diversity.

## Statistical analysis

All the enzyme activity experiments were performed thrice, each set in duplicates. The data are shown as averages with standard deviations. The Student’s T-test for statistical significance was calculated using GraphPad Prism 8.0.2 software.

## CRediT authorship contribution statement

**Simran Sharma:** Performed all the wet lab experiments as well as some of the docking studies.

**Vijayan Ramachandran-** Performed the *in silico* screening of the potential inhibitors.

**Rohini Muthuswami-** Conceptualised the project, brought in the funds, advised on experiments and read/edited the manuscript.

**Samudrala Gourinath-** Conceptualised the project, brought in the funds, advised on experiments and read/edited the manuscript.

**Sneha Sudha Komath-** Conceptualised and supervised the project and brought in the funds required; wrote the overall framework of the manuscript and first draft.

All authors have read, edited/ commented and agreed with the final version of the manuscript.

## Acknowledgements

Parts of this work were funded by grants to S.S.K. from Department of Biotechnology (DBT) India BT/PR53604/BMS/85/165/2024, BT/PR51192/MED/29/1662/2023 (as co-PI) and the MHRD-STARS grant to RM, SSK, SG (STARS/APR2019/BS/224/FS). Facilities and funding support by ANRF-PAIR (ANRF/PAIR/2025/000029/PAIR) and DBT-BUILDER (BT/INF/22/SP45382/2022) are also gratefully acknowledged. SS was supported by DBT Junior and senior research fellowships. The facilities at the Central Instrumentation Facility (CIF), School of Life Sciences (SLS), JNU have also been supported by grants from UGC-CAS, UGC-RNW, and DST-FIST (II).

## Data availability statement

The main body of data, including analyses and images, is available in the article or its Supporting Information. Sources of data are available from the corresponding authors upon reasonable request.

## Abbreviations

AcCoA: acetyl-CoA
BLI: biolayer interferometry
DTT: dithiothreitol
GFC: gel filtration chromatography
HAT: histone acetyltransferase
IC_50_: concentration for 50% inhibtion
k_cat_: turnover number
K_D_: dissociation constant
K_i_: inhibition constant
K_M_: Michaelis-Menten constant
V_max_: maximum velocity/reaction rate

## Supporting Information

Figure S1. Confirmation of *CaRTT109*, *CaRTT109Δ_Loop_*, *CaVPS75*, *CaVPS75Δ_C_*, and *CaASF1* cloned in bacterial expression vectors.

**Figure S2. Calibration curve for the gel filtration chromatography columns.**

**Figure S3. Gel filtration analysis of CaRtt109 co-incubated with its chaperones. SDS-PAGE for the GFC profiles shown in** **Figure 5**.

**Figure S4. Cloning, expression, purification and HAT activity of ScGcn5.**

**Figure S5. Real-time spectra of HAT activity of CaRtt109 without and with its chaperones.**

