## Supporting information for "Purification and characterization of recombinant Rtt109, a fungus-specific histone acetyltransferase, from *Candida albicans*"

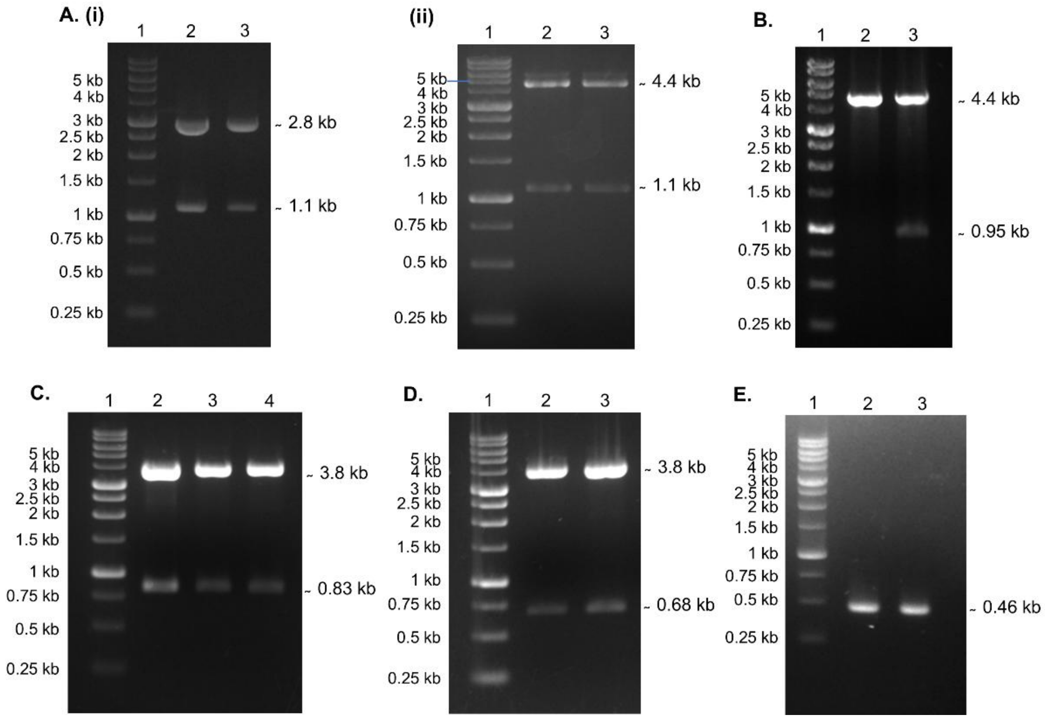


**Figure S1. Confirmation of *CaRTT109*, *CaRTT109*$\Delta_{Loop}$, *CaVPS75*, *CaVPS75*$\Delta_{C}$, and *CaASF1* cloned in bacterial expression vectors. A. Confirmation of *CaRTT109* cloned in TA and pCold I vector**. **(i)** Cloning confirmation of *CaRTT109* in TA vector. Lane 1: 1 kb DNA ladder, Lane 2, 3: positive clones showing TA vector of ~2.8 kb and *CaRTT109* gene of ~1.1 kb upon double restriction digestion. **(ii)** Cloning confirmation of *CaRTT109* in pCold I vector, Lane 1: 1 kb DNA ladder, Lane 2, 3: positive clones showing pCold I vector of ~4.4 kb and *CaRTT109* gene of ~1.1 kb upon double restriction digestion. **B. Confirmation of *CaRTT109*$\Delta_{Loop}$ cloned in pCold I.** Lane 1: 1 kb DNA ladder, Lane 2: Negative clone. Lane 3: Positive clone showing pCold I vector of ~4.4 kb and *CaRTT109*$\Delta_{Loop}$ of ~0.95 kb upon double restriction digestion. **C. Confirmation of *CaVPS75* cloned in pRSFDuet1.** Lane 1: 1 kb DNA ladder, Lane 2, 3, 4: positive clones showing pRSFDuet1 vector (~ 3.8 kb) and *CaVPS75* (~ 0.83 kb) upon double restriction digestion. **D. Confirmation of *CaVPS75*$\Delta_{C}$ cloned in pRSFDuet1.** Lane 1: 1 kb DNA ladder, Lane 2,3: positive clones showing pRSFDuet1 (~3.8 kb) and *CaVPS75*$\Delta_{C}$ (~0.68 kb) upon double restriction digestion. **E. Confirmation of *CaASF1* cloned in pCold I vector.** Lane 1: 1 kb DNA ladder, Lane 2, 3: positive clones confirmed by plasmid PCR showing 0.46 kb *CaASF1* gene.

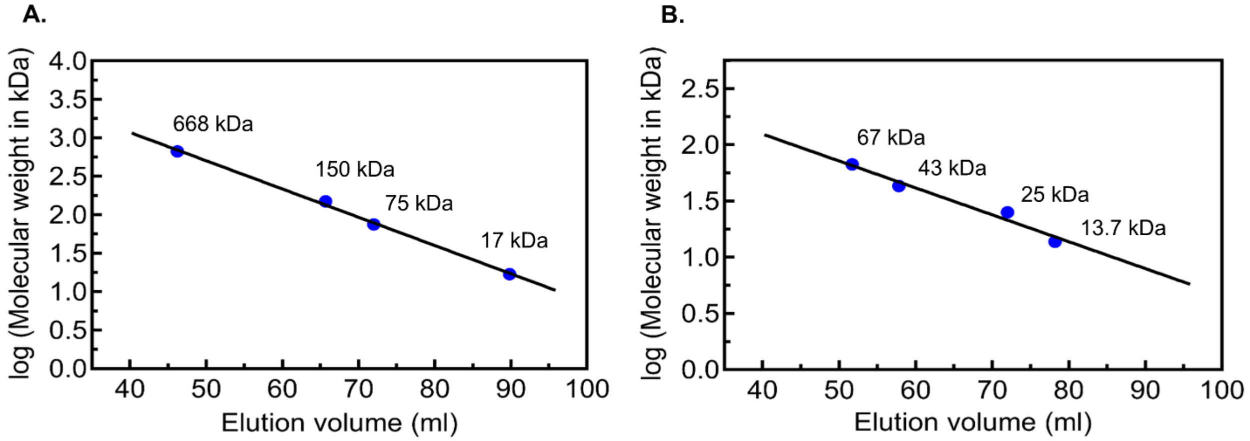


**Figure S2. Calibration curve for the gel filtration chromatography columns. A. Calibration curve for HiLoad 16/60 Superdex 200 column.** The following protein standards were employed: myoglobin (17 kDa), conalbumin (75 kDa), alcohol dehydrogenase (150 kDa) and thyroglobulin (668 kDa). **B. Calibration curve of HiLoad 16/60 Superdex 75 column.** The following protein standards were employed: Ribonuclease A (13.7 kDa), chymotrypsin (25 kDa), ovalbumin (43 kDa) and bovine serum albumin (67 kDa).

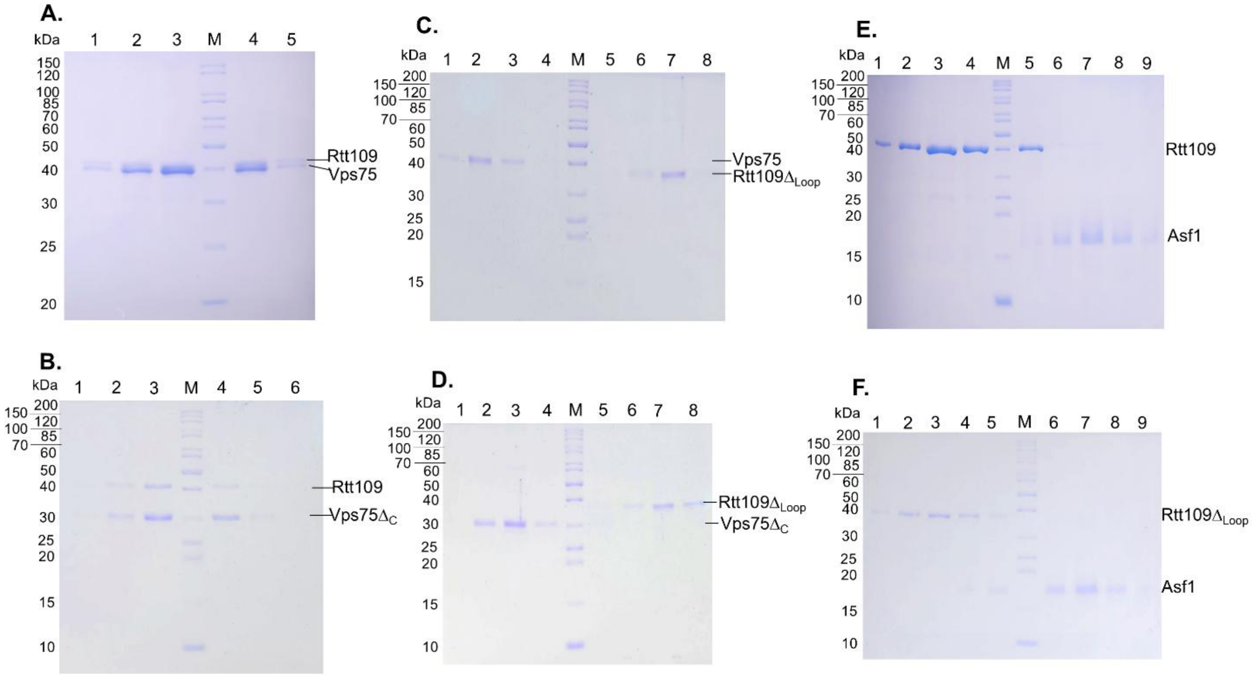


**Figure S3. Gel filtration analysis of CaRtt109 co-incubated with its chaperones. SDS-PAGE for the GFC profiles shown in Figure 5. A. Co-elution of CaRtt109 with CaVps75**. CaRtt109 and CaVps75 were separately purified using Ni$^{+2}$-NTA affinity chromatography followed by GFC. CaRtt109 (10 µM) was incubated with CaVps75$\Delta_{C}$ (40 µM) at 4 °C overnight, and GFC was performed. The eluted fractions (2 ml each) were analysed using 10% SDS-PAGE. Lane M: Molecular weight markers; Lane1-5: eluted fractions at 60 ml, 64 ml, 68 ml, 72 ml, and 76 ml, respectively. **B. Co-elution of CaRtt109 with CaVps75$\Delta_{C}$.** CaRtt109 and CaVps75$\Delta_{C}$ were separately purified using Ni$^{+2}$-NTA affinity chromatography followed by GFC. CaRtt109 (10 µM) was incubated with CaVps75$\Delta_{C}$ (40 µM) at 4 °C overnight, and GFC was performed. The eluted fractions (2 ml each) were analysed using 12% SDS-PAGE. Lane M: Molecular weight markers; Lane1-6: eluted fraction at 58 ml, 62 ml, 66 ml, 68 ml, 72 ml, and 76 ml, respectively. **C. Co-elution of CaRtt109$\Delta_{Loop}$ with CaVps75.** CaRtt109$\Delta_{Loop}$ and CaVps75 were separately purified using Ni$^{+2}$-NTA affinity chromatography followed by GFC. CaRtt109$\Delta_{Loop}$ (10 µM) was incubated with CaVps75 (40 µM) at 4 °C overnight, and GFC was performed. The eluted fractions (2 ml each) were analysed using 12% SDS-PAGE. Lane1-8: eluted fraction at 58 ml, 62 ml, 66 ml, 70 ml, and 74 ml, 80 ml, 84 ml, and 88 ml, respectively. **D. Co-elution of CaRtt109$\Delta_{Loop}$ with CaVps75$\Delta_{C}$.** CaRtt109$\Delta_{Loop}$ and CaVps75$\Delta_{C}$ were separately purified using Ni$^{+2}$-NTA affinity chromatography followed by GFC. CaRtt109$\Delta_{Loop}$ (10 µM) was incubated with CaVps75$\Delta_{C}$ (40 μM) at 4 °C overnight, and GFC was performed. The eluted fractions (2 ml each) were analysed using 12% SDS-PAGE. Lane M: Molecular weight markers; Lane1-8: fraction eluted at 58 ml, 62 ml, 66 ml, 70 ml, and 74 ml, 80 ml, 84 ml, and 88 ml, respectively. **E. Co-elution of CaRtt109 with CaAsf1.** CaRtt109 and CaAsf1 were separately purified using $Ni^{+2}$-NTA affinity chromatography followed by GFC. CaRtt109 (10 μM) was incubated with CaAsf1 (20 μM) at 4 °C overnight, and GFC was performed. The eluted fractions (2 ml each) were analysed using 12% SDS-PAGE. Lane M: Molecular weight markers; Lane1-9: eluted fraction at 56 ml, 58 ml, 60 ml, 62 ml, 64 ml, 66 ml, 68 ml, 72 ml, and 76 ml, respectively. **F. Co-elution of CaRtt109$\Delta_{Loop}$ with CaAsf1.** CaRtt109$\Delta_{Loop}$ and CaAsf1 were separately purified using $Ni^{+2}$-NTA affinity chromatography followed by GFC. CaRtt109$\Delta_{Loop}$ (10 μM) was incubated with CaAsf1 (20 μM) at 4 °C overnight, and GFC was performed. The eluted fractions (2 ml each) were analysed using 12% SDS-PAGE. Lane M: Molecular weight markers; Lane1-9: eluted fraction at 58 ml, 62 ml, 64 ml, 66 ml, 68 ml, 70 ml, 72 ml, 74 ml, and 76 ml, respectively.

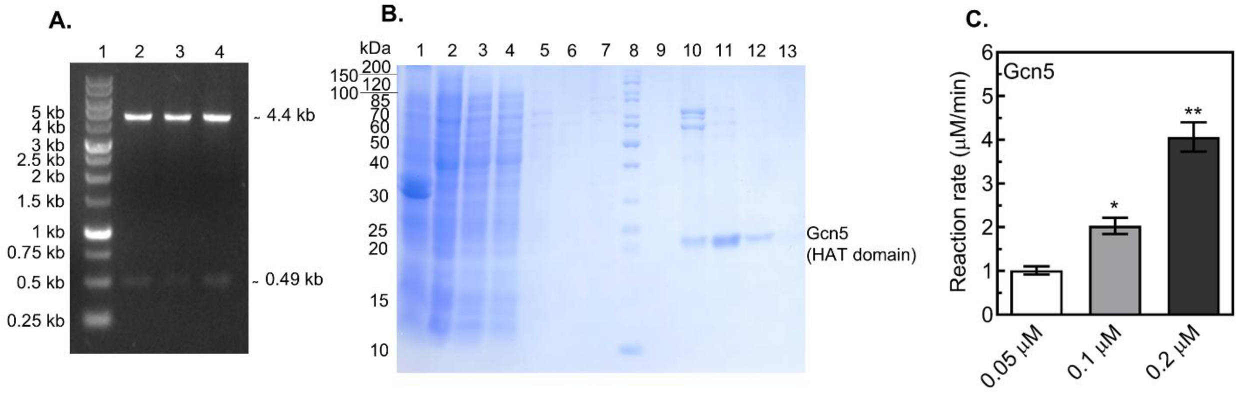


**Figure S4. Cloning, expression, purification and HAT activity of ScGcn5. A. Cloning confirmation of pColdI-*ScGCN5*.** Lane 1: 1 kb DNA ladder, Lane 2, 3: Positive clones showing pCold I vector of ~4.4 kb and *ScGCN5* gene segment of ~0.49 kb upon double restriction digestion. **B. SDS-PAGE analysis of the expression and purification of ScGcn5.** *E. coli* Rosetta(DE3) cells transformed with the respective recombinant plasmid were grown until $OD_{600nm}$ ~0.6 and protein expression induced using 0.5 mM IPTG at 16 ºC for 16 h. The cells were harvested and lysed in 50 mM HEPES-KOH, pH 7.5, containing 150 mM NaCl, 0.5 mM DTT, 5% glycerol, and 1 mM PMSF. Samples from different steps of the $Ni^{+2}$-NTA affinity purification were analyzed using 12% SDS-PAGE. Lanes 1 and 2: Pellet and supernatant fractions, respectively, obtained after centrifugation of the cell lysate; Lane 3: Flow-through fraction from the $Ni^{+2}$-NTA column, showing the unbound proteins; Lanes 4-5: wash fractions with buffer containing 300 mM NaCl and 500 mM NaCl, respectively; Lane 6 and 7: wash fraction with buffer containing 150 mM NaCl with 5 mM and 10 mM imidazole, respectively ; Lane M: Molecular weight markers; Lanes 9-13: protein fractions (2 ml each) eluted in the presence of 200 mM imidazole. **C. HAT activity of ScGcn5.** The activity assay was performed with ScGcn5 HAT at different concentrations (0 µM, 0.05 µM, 0.1 µM, and 0.2 µM) using pyruvate dehydrogenase coupled assay. The reactions were performed at 100 µM H3 peptide and 50 µM AcCoA. The average initial reaction rates were plotted with standard deviation from three different experiments. The symbol * and ** show statistical significance with p-value <0.05 and <0.01, respectively.

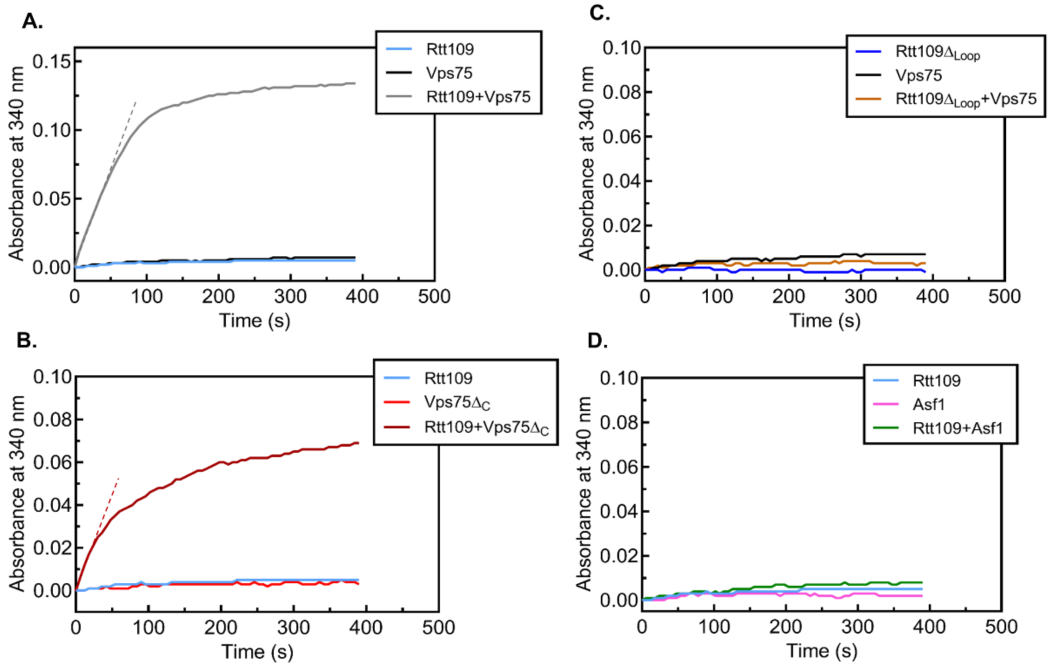


**Figure S5.** Real-time spectra of HAT activity of CaRtt109 without and with its chaperones. The HAT activity of CaRtt109 on H3 peptide was monitored spectrophotometrically using a pyruvate dehydrogenase coupled assay. The resulting NADH formed, which is linked directly to the amount of acetylation, is continuously measured at 340 nm with time. The assay was performed in the presence of 200 μM H3 peptide and 50 μM AcCoA. In A. and B. the slope is represented by dashed line. **A. Activity of CaRtt109 in the presence of CaVps75.** Representative plot for the continuous change in absorbance with time in the presence of either 0.5 μM CaRtt109, 2 μM CaVps75 or both. **B. Activity of CaRtt109 in the presence of CaVps75ΔC.** Representative plot for the continuous change in absorbance with time in the presence of either 0.5 μM CaRtt109, or 2 μM CaVps75ΔC, or both. **C. Activity of CaRtt109ΔLoop in the presence of CaVps75.** Representative plot for the continuous change in absorbance with time in the presence of either 0.5 μM CaRtt109ΔLoop, 2 μM CaVps75, or both. **D. Activity of CaRtt109 in the presence of CaAsf1.** Representative plot for the continuous change in absorbance with time in the presence of either 0.5 μM CaRtt109, or 2 μM CaAsf1, or both.
